# Insights for Estimating Animal Movement Step Selection Functions

**DOI:** 10.64898/2026.08.29.748012

**Authors:** Phillip Koshute, William F. Fagan

## Abstract

Ecologists remotely track movement steps of animals (e.g., via global positioning systems) and use step selection functions to study the effect of environmental factors upon their movement decisions. Constructing such functions requires pairing each observed step with some number of unobserved but feasible comparison steps. Larger numbers of comparison steps generally yield better estimates but also incur potentially challenging computational demands. Thus, it is important to determine an appropriate number of comparison steps. No established guidance exists for this decision. Here, we use simulated tracks to assess how many comparison steps are needed, fitting each set of steps to a conditional logistic regression model. We monitor errors in estimated effects for several classes of tracks, identifying the number of comparison steps for which mean relative absolute error in estimated effects is consistently low. By this criterion, 32 comparison steps per observed step are needed for our primary class of simulated tracks. Tracks in more homogeneous landscapes, tracks with shorter mean step lengths, or shorter tracks generally require more comparison steps (ranging from 64 – 128 per observed step) to achieve the same level of accuracy. Longer tracks generally require fewer comparison steps (16 per observed step). These results clearly demonstrate that the number of comparison steps influences how well step selection functions estimate covariate effects and provides initial direction in a research area that currently lacks quantitative guidance. Movement ecologists should take care when selecting the number of comparison steps paired with each observed step because those decisions matter.

## 1. Background

Ecologists study the movement of animals in order to better understand animals’ relationships with each other and with their surroundings. This research aids scholarly and conservation efforts alike (Costa-Pereira et al. 2022; Katzner and Arlettaz 2020). By understanding the factors that motivate animal movement, accommodations may be made to counteract naturally occurring or manmade landscape changes. For instance, movement data from Florida panthers (*Puma concolor coryi*) were used to determine optimal locations for crossings to accommodate panthers’ continued freedom of movement (Loraamm and Downs 2016).

In the last half-century, spatial ecological research has been aided by increased availability and capability of remote tracking technology. In such studies, animals are outfitted with collars or backpacks that include global positioning system (GPS) units that periodically record and transmit time-stamped data on the animals’ locations (Kays et al. 2015). Considered together, the recorded locations (“fixes”) for a given animal form that animal’s “track” over the period of study. A pair of consecutive fixes comprises a “step” from one recorded location to the next recorded location.

These tracks provide a discrete approximation of an animal’s path. However, on their own, they do not indicate why the animal has followed that path. In practice, animals’ movements are driven by various factors including resource abundance, shelter availability, proximity of predators, and presence of human infrastructure.

Step selection functions (SSFs) provide a framework for studying the factors that drive animals’ movements. This technique was introduced by Fortin et al. (2005) as a way of studying the behavior of elk in the presence of wolves. By comparing the factors at feasible but unobserved steps with the conditions at observed steps (OSs), SSFs infer the factors that motivate animals’ selected steps. These unobserved steps included for fitting an SSF are often called available steps, random steps, or “comparison steps” (CSs). The set consisting of a single OS and all corresponding CSs comprise a “stratum” of steps.

SSFs have been used by dozens of studies to investigate how animals’ environments affect their movement decisions (e.g., Fortin et al. 2005; Barry et al. 2020; Eisaguirre et al. 2020). In some cases, these investigations contribute to practical decisions related to conservation (Loraamm and Downs 2016). See Thurfjell et al. (2014) for a broad overview of SSFs. For most of these studies, the strata of OSs and their corresponding CSs are used to fit a conditional logistic regression (CLR) model (Hosmer et al. 2013). Similar to (unconditional) logistic regression, this approach models the probability of a given outcome as a function of a weighted sum of covariates. For SSFs, the outcome is whether a given step will be observed. Unlike logistic regression, CLR involves data that are explicitly split into strata and a separate likelihood function is computed for each stratum.

Within SSFs, fitted CLR models and their estimated covariate coefficients (“effects”) provide quantitative results on the factors affecting animals’ movement. Typical statistical software outputs each covariate’s estimated effect, standard error, z-score, p-value, and significance level. Researchers may interpret these results in several different ways: directly assessing the estimated effects, comparing the estimated effects (in terms of both signs and relative magnitudes), or using estimates’ p-values to assess the statistical significance of the covariates. Researchers may also fit models to different subsets of covariates and identify the subset that corresponds to the smallest value of the Akaike information criteria (AIC) or other model selection criteria.

The selection of CSs affects the extent to which an SSF is able to accurately fit the CLR model and ultimately infer the factors motivating an animals’ movements within its track. A CS is characterized by a step length and turning angle. Typically, these values are drawn from distributions stemming from animals’ OSs (Thurfjell et al. 2014).

Researchers must also determine the number of CSs to be paired with each OS. This decision is less straightforward. Having more CSs (larger *M*) enables a better approximation of the landscape within which the animal has made their step decisions (Fieberg et al. 2021). However, using more CSs also increases runtime for the SSF analysis and computer storage requirements. Modern GPS units now routinely yield movement tracks exceeding 10^6^ fixes (Kays et al. 2015). Hence, depending on the number of OSs within an animal’s track, the computational requirements for some numbers of CSs can be prohibitive. On the other hand, having fewer CSs risks inaccurate effect estimates and potentially incorrect conclusions about the extent to which particular environmental factors affect animals’ movement decisions (if at all).

Currently, no clear guidance exists as to how many steps are required to achieve a certain level of accuracy in inferring the effects of the factors that motivate an animal’s movement decisions. In published studies involving SSFs, this number of CSs paired with each OS ranges from 1 to 200. In many of these studies, no justification is given for the number of CSs.

Several studies have offered insights on appropriate numbers of CSs for SSFs but provide limited direct guidance. Observing that many CSs can yield “excessive” data sets and pose “computational limitations,” Thurfjell et al. (2014) suggested that few or even one CS per OS could suffice. In contrast, Fieberg et al. (2021) advocated for using many CSs per OS (“the more points the merrier”), refraining from recommending a particular number. In their original paper, Fortin et al. (2005) used 200 CSs per OS, striving to include rare values of one covariate of interest in each stratum. Michelot et al. (2024) discussed how different approaches for how the CSs are sampled, i.e., the underlying distribution of sampled CSs. Giving more weight to likely CSs (e.g., according to an approximate movement kernel) generally reduces the number of CSs needed.

Other studies have discussed the number of available locations that should be sampled within the similar technique of resource selection functions (RSFs) (Manly et al. 2002). RSFs consider locations visited by an animal (at any time) rather than particular steps (taken at a particular time), comparing used locations with other available locations. Warton and Shepherd (2010) and Fithian and Hastie (2012) provided theoretical justification for using logistic regression models, as long as a “sufficiently large” number of available locations are sampled. Northrup et al. (2013) found that RSF effect estimates stabilized with approximately 10000 available locations (not for each used location, but in total). Street et al. (2021) proposed a helpful analytic solution for the numbers of individual animals (when multiple animals are studied together) and the number of used locations per individual needed to achieve specified precision for RSF effect estimates; however, they do not incorporate the number of available locations in their solution, instead sampling one available location per used location.

Within our study, we aim to shed further light on what might determine an appropriate number of CSs to be paired with each OS when fitting an SSF. We propose a concrete process for rigorously assessing how many CSs are needed for a given track. As a way to explore a variety of well-defined scenarios, we leverage simulation and apply our process to simulated covariate landscapes and simulated tracks. We focus on tracks with a given set of characteristics and extend our analysis to tracks with different characteristics. This analysis yields recommendations for numbers of CSs for certain classes of tracks. In laying out our approach, we likewise aim to pave the way for a general strategy that would be applicable to an SSF for any animal track.

## 2. Methods

To address the question of how many CSs are sufficient for a given animal track’s SSF, we simulate dozens of tracks across multiple simulated landscapes, specifying the effects of different environmental covariates on the movement process. For each track, we sample many replicate sets of different numbers of CSs for each OS. We fit a CLR model to each replicate set, yielding hundreds of fitted models for each track. From each model’s results, we study the estimated effects of the covariates, assessing how these estimates change as the number of CSs increases. Ultimately, we search for a number of CSs that enables the effects of covariates to be estimated with satisfactory accuracy and limited variance. For a given track, it is this number of CSs that a researcher needs to pair with each OS to confidently leverage SSFs for inferring effects of environmental factors on animal movement. We have conducted all analyses and constructed all plots using R (R Core Team 2024).

### 2.1. Step Selection Functions

SSFs presume that animals have consistent patterns that generally guide their movement decisions and that measured covariates suffice to approximate these patterns. As discussed above, SSFs seek to distinguish between the characteristics of OSs (shown as a blue solid arrow in Fig. 1) with the characteristics of unobserved but feasible CSs (shown as red dashed arrows). Roughly, if the distribution of the values of a given covariate is similar among both OSs and CSs, SSFs will indicate that that covariate does not strongly affect movement decisions. However, if the covariate’s values are consistently higher or lower at OSs (relative to CSs and controlling for the values of other covariates), SSFs will highlight the effect of that covariate upon the animal’s movement decision. For those unfamiliar with SSFs, we provide further details in the Supplemental Material (Appendix A), recapping applicable notation, motivating concepts, modeling decisions, and possible analysis objectives.

**Fig. 1.**
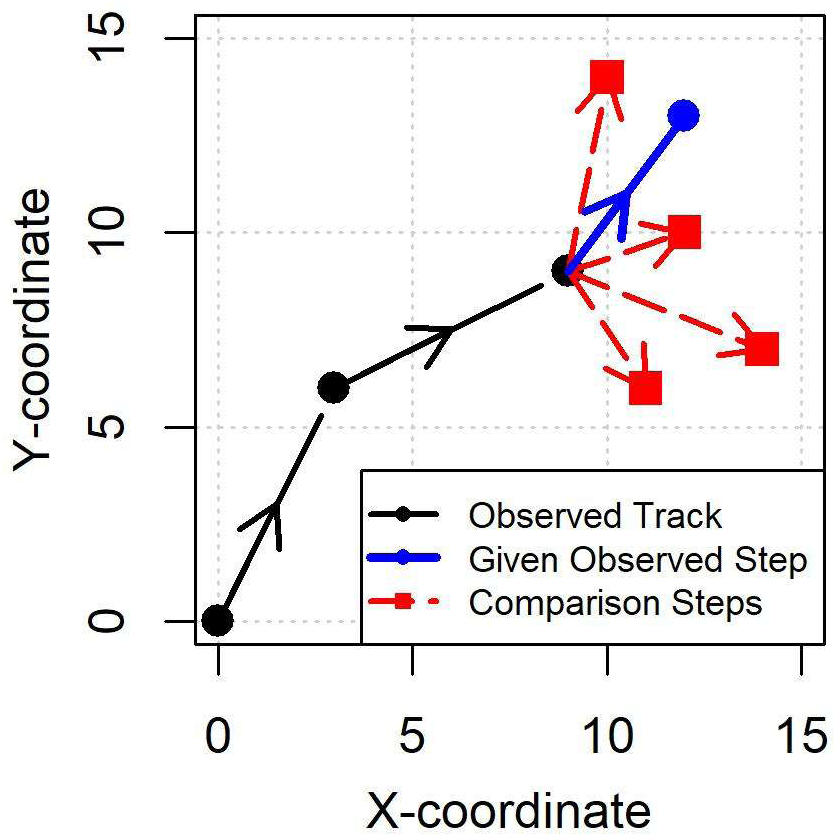
Steps within an Example Animal Track

Following Fortin et al. (2005), we estimate the effects ***β*** for SSF covariates within a CLR model, pairing each OS with *M* CSs. The CSs originating from a given location are drawn from a distribution *F* of feasible alternative steps (characterized by parameters such as mean step length, which are represented with γ).

The CLR model involves the following probability, which is conditioned upon an animal’s current location ***x_t_*** and is a function *f* of covariates ***z^m^_t+1_*** that reflect environmental factors or other characteristics for a given OS or CS that ends at ***x^m^_t+1_***:

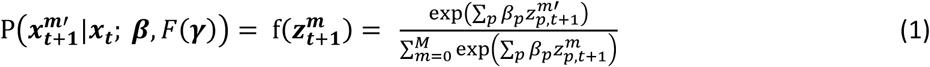

Computationally, the effects ***β*** are estimated by considering the product of the likelihood equations for each stratum (Hosmer et al. 2013). In practice, these effects are often estimated with the *clogit* function from the *survival* package (Therneau 2015) in R. As mentioned above, the number of CSs drawn from *F*(γ) and paired with each OS has varied across studies. Determining a minimum number of CSs needed for accurately inferring covariates’ effects is the focus of this analysis.

Because animals tend to continue in the same direction from one step to another, locations close to each other within a given animal’s track are often correlated (Fleming et al. 2014). Such autocorrelation is particularly likely when the animal’s locations are transmitted with high frequency. One technique to account for autocorrelation is to split the data into independent clusters of autocorrelated locations (Prima et al. 2017). This enables the use of general estimating equations (GEE) (Hardin and Hilbe 2002) to estimate the covariance matrix of the effects ***β***, while accounting for the autocorrelation. Nevertheless, most existing SSF studies do not follow these recommendations, instead estimating the variance of each effect directly from the fitted CLR model. To be consistent with this common practice, we similarly use the estimated effects’ standard errors directly reported for the fitted model (i.e., from the *clogit* function). We leave an analysis of the impact of using GEE upon the number of CSs needed as future work.

### 2.2. Simulation Details

We fit CLR models to simulated tracks within simulated landscapes with different numbers of CSs and many replicates for each number of CSs. We repeat this analysis with different landscapes (characterized by smoothing window size) and different track characteristics (mean step length, track length) in order to study how such characteristics influence the “convergence” of SSF models with increasing numbers of CSs. (We discuss this convergence in the next subsection.) At each location in the landscape, we simulate a set of covariates, corresponding to different resources or other measurable factors.

To simulate the landscape of values for a given covariate, we set up a square grid of cells. For each cell, we sample an initial value from a normal distribution with mean zero and standard deviation one. To obtain different amounts of correlation within the grid, we use a smoothing window with width 2*W* + 1. For instance, when *W* = 1, the smoothed value of one cell is the mean of the initial values of that cell and the eight nearest neighbor cells. Larger values of *W* achieve greater spatial correlation.

We simulate separate grids for each covariate and do not model any correlation between the grids of different covariates. Having correlation between covariates could be added within future work. We simulated grids for three covariates, two of which actually influence the movement tracks and the third of which is a ‘noise covariate’ representing a landscape feature that a researcher might reasonably hypothesize to influence movements but which ultimately does not. See the Supplemental Material (Appendix B) for our analysis of the impact of the noise covariate.

Given a simulated landscape, we simulate tracks by successively carrying out weighted sampling of 10000 candidate steps near the track’s current location. We determine the candidate steps according to randomly sampled step lengths and relative turning angles. (A “relative” turning angle of a given step measures the angular difference between that step and the straight-line extension of the previous step.) We draw step lengths *L* from an exponential distribution with mean *μ_L_*, assessing different values of *μ_L_*. We do not specify a unit for step length; instead, step length can be interpreted according to the spatial resolution of the covariate landscape. For instance, a step length of one corresponds to the width of a single grid within the covariate landscape. We assume that all covariates have the same spatial resolution. Similarly, we draw relative turning angles *A* from a von Mises distribution with concentration parameter *κ_A_* = 3. Correlation between the step length and relative turning angle could be studied for particular animals, but we do not model or analyze such correlation in this study.

For sampling the candidate steps, we compute the weights of each candidate step from the covariate values at the step destination, e.g., rather than computing an average value along the entire (linear) step. Focusing on the values at a step destination is a common practice for SSFs (Thurfjell et al. 2014). For each covariate, we specify effects that determine the extent to which that covariate affects the sampling weight of each candidate step (and ultimately its contribution to step selection decisions): *β_1_* = 4 as the effect of the first covariate and *β_2_* = 2 as the effect of the second covariate. (Due to sampling noise, finite simulated tracks generally do not exactly reflect these specified effects. See Appendix C in the Supplemental Material for more details on this phenomenon.) The third covariate does not influence simulated movement decisions; effectively, *β_3_* = 0. We focus our analysis on models without *β_3_* but also consider the impact of including *β_3_* because, in practice, researchers may hypothesize that environmental factors are important determinants of movement when in reality they are not. (See Appendix B for results with models involving *β_3_*. We found that increasing the number of CSs does not always enable a model to distinguish a noise factor such as *β_3_*.)

Covariates’ contributions *λ* for the *m*th candidate step (ending at location [inine]) are summarized by the resource-based selection probability in Equations 2 and 3. We have *P* = 2 or 3 total covariates.

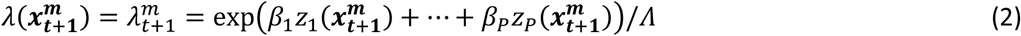

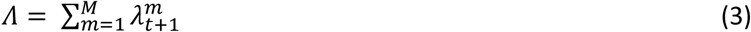

Ultimately, we set the weight of each candidate step as *λ^m^_t+1_* (see Equation 2). Candidate steps with larger weights are more likely but not guaranteed to be sampled.

We continue to iteratively simulate steps according to this process until we have simulated a track with the specified track length *N*. The resulting track has *N*+1 fixes and *N* OSs. Because a prior step (involving two prior locations) is required to compute relative turning angle, we use *N*-1 OSs for each model.

We refer to tracks with a common set of simulation characteristics as a “class” of tracks. Within this study, we focus on tracks with length *N* = 1000 and specified mean step length *μ_L_* = 4 within landscapes with smoothing window size *W* = 2, referring to this class as the “base” class. We also consider classes of tracks with different lengths (*N* = 500 and *N* = 2000), different specified mean step lengths (*μ_L_* = 2 and *μ_L_* = 8), and different landscape smoothing window sizes (*W* = 1 and *W* = 4). Table 1 summarizes the characteristics for each class of tracks. To account for random variation in particular landscapes and tracks, we simulate five tracks for each of five covariate landscapes, totaling 25 tracks for each class and 175 tracks overall. Additional detail on our simulation process is included in the Supplemental Material (Appendix D).

**Table 1:**
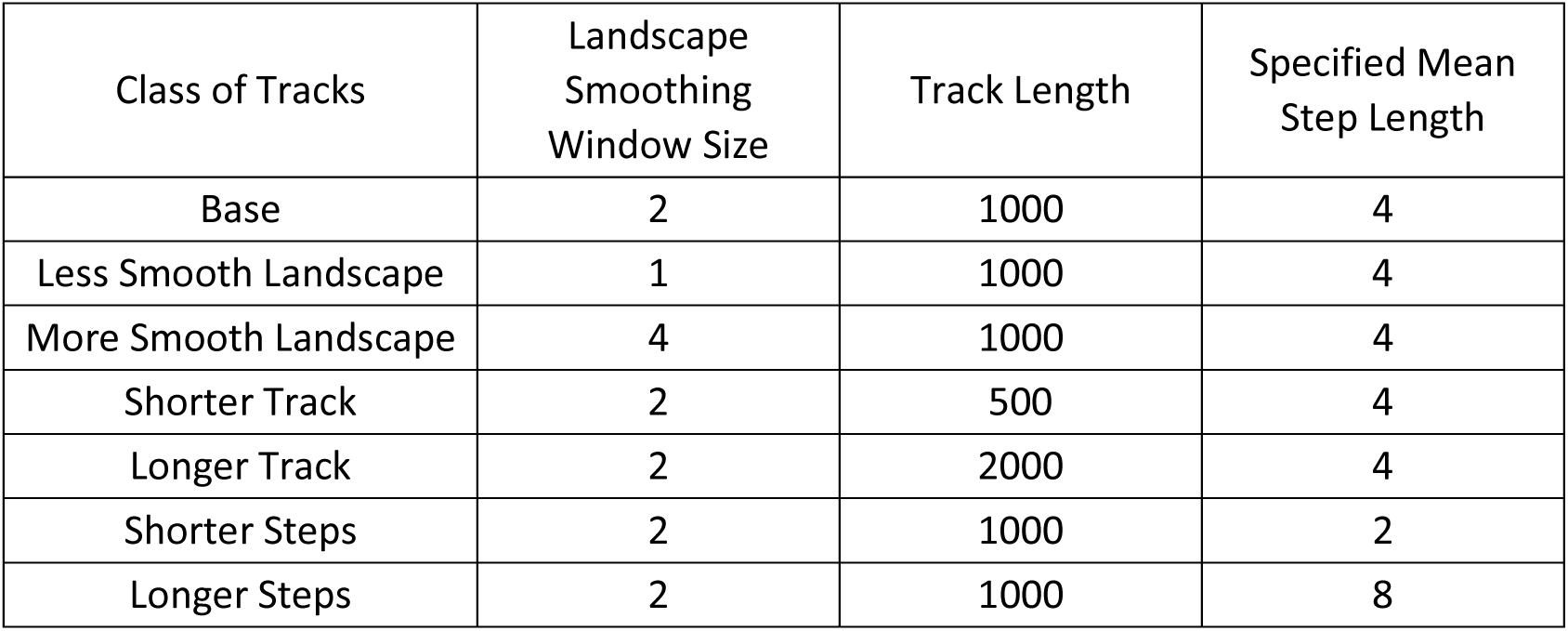
Summary of Simulated Classes of Tracks.

### 2.3. Analysis Approach

To determine the number of CSs needed for each track, we study how fitted models and their corresponding effects change as the number of CSs (*M*) changes. Additionally, we establish a concept of “convergence” for models and their estimates. Using this concept, we identify the value of *M* at which the models and estimates have converged. It is this *M* that we regard as the number of CSs needed.

Studies with SSFs can have different objectives, including estimating effects of CLR covariates and selecting models with the optimal subset of covariates. (See Table S-1 in the Supplemental Material for a recap of these objectives.) We have confirmed that greater *M* enables more precise and accurate effect estimation but not necessarily better model selection. For instance, we have found some tracks for which greater *M* yields poorer model selection. This phenomenon of selecting models with “false positive” significant effects is discussed by Sutherland et al. (2023). (See also Appendix B.) In light of these inconsistencies with SSF model selection, we focus on the influence of *M* on estimating effects for CLR models. In particular, we seek to identify the value of *M* required to have a specified level of accuracy in effect estimation with a specified level of consistency.

In order to fit the CLR models, we sample *M* CSs for each OS. Our approach for sampling CSs is similar to how we sample candidate OSs while simulating tracks, except that we do not include covariate-based weighted sampling. Rather, following the findings of Michelot et al. (2024), we use Monte Carlo sampling according to an approximate movement kernel (a special case of “importance sampling”), determining CSs according to randomly sampled step lengths and relative turning angles that originate from fixes along the observed track. As in Section 2.2, we draw step lengths *L* from an exponential distribution with mean *μ_L_* and relative turning angles *A* from a von Mises distribution with concentration parameter *κ_A_*. We fit both *μ_L_* and *κ_A_* to the observed track, fitting *μ_L_* with the *fitdistr* function from the *MASS* package and fitting *κ_A_* with the *lm.circular* function from the *circular* package. These fitted values are typically similar to but not necessarily the same as the specified values for these parameters. (See Appendix E in the Supplemental Material for an exploration of the underlying causes of this discrepancy.) This sampling approach yields *M*+1 steps per stratum (one OS and *M* CSs) that are available for fitting a CLR model.

Following Avgar et al. (2016), we model an individual animals’ step selection decisions according to steps’ resource-based covariate values, as well as their lengths and the cosines of their relative turning angles. This “integrated step selection” simultaneously estimates resource-based covariate effects and updates the estimates for the parameters for step length and turning angle distributions. For the *m*th possible step originating from ***x_t_***, we denote the step length as *l^m^_t+1_* and the relative turning angle as α*^m^_t+1_*; the *p*th covariate for ***x^m^_t_*** is *Z^m^_t,p_*. Using this approach, we essentially regard the step length and the cosine of the relative turning angle as additional covariates.

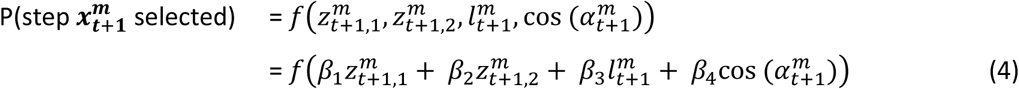

For each track, we sample thirteen different values of *M* for each OS, corresponding to powers of two: 1, 2, 4, …, 4096. To robustly assess the trend as *M* changes, we sample 100 replicates with each *M*. Each replicate has a different set of CSs for each OS. This amounts to 1300 replicate sets of CSs and fitted models for each track. Using these results, we assess model performance with increasing *M*, identifying how many CSs are necessary to reach specified levels of accuracy in covariate effect estimation.

We assume that models fitted with very many CSs (for us, *M* = 4096) yield reasonably accurate estimates of the true simulated effects ***β\****. This assumption accords with Michelot et al. (2024), who used “asymptotic estimates” obtained with very many CSs (5000) per OS as approximations of the true effects within a track. For the *i*th covariate for a given track, we use the mean of all estimates β̂_*i*_ with *M* = 4096 as β_*i*_^*^. Likewise, for each covariate *β*_i_, we compute a relative error *RE_i_* = (β̂_i_ − β^*^_*i*_)/β^*^_*i*_. Again, following Michelot et al. (2024), we anticipate that models fitted with more CSs will have lower relative absolute errors (RAEs). We want to assess what *M* is needed in order to have satisfactorily low RAEs. We denote this threshold as *ε_1_*. Within this study, we set *ε_1_* = 0.05, i.e., setting 5% as a tolerable level of RAE.

For each track *j*, we have 1300 RAEs (one for each fitted model, with 100 models fitted for each of the 13 values of *M*). To determine whether a given value of *M* suffices, we compute *q^i^_2,M_*, which is the 100*(1-*ε_2_*)^th^ quantile of RAEs for covariate *i* with *M* CSs. Within this study, we set *ε_2_* = 0.1, i.e., setting the maximum acceptable rate at which the tolerable RAE level is exceeded as 10%. Thus, among models fitted with *M* CSs for each OS in that track, no more than 100\**ε_2_* = 100*0.1 = 10 models (out of 100) will have RAEs that exceed *q^i^_2,M_*. Accordingly, we can plot *M* versus *q^i^_2,M_*; see plots C and D within Fig. 2 as examples.

**Fig. 2.**
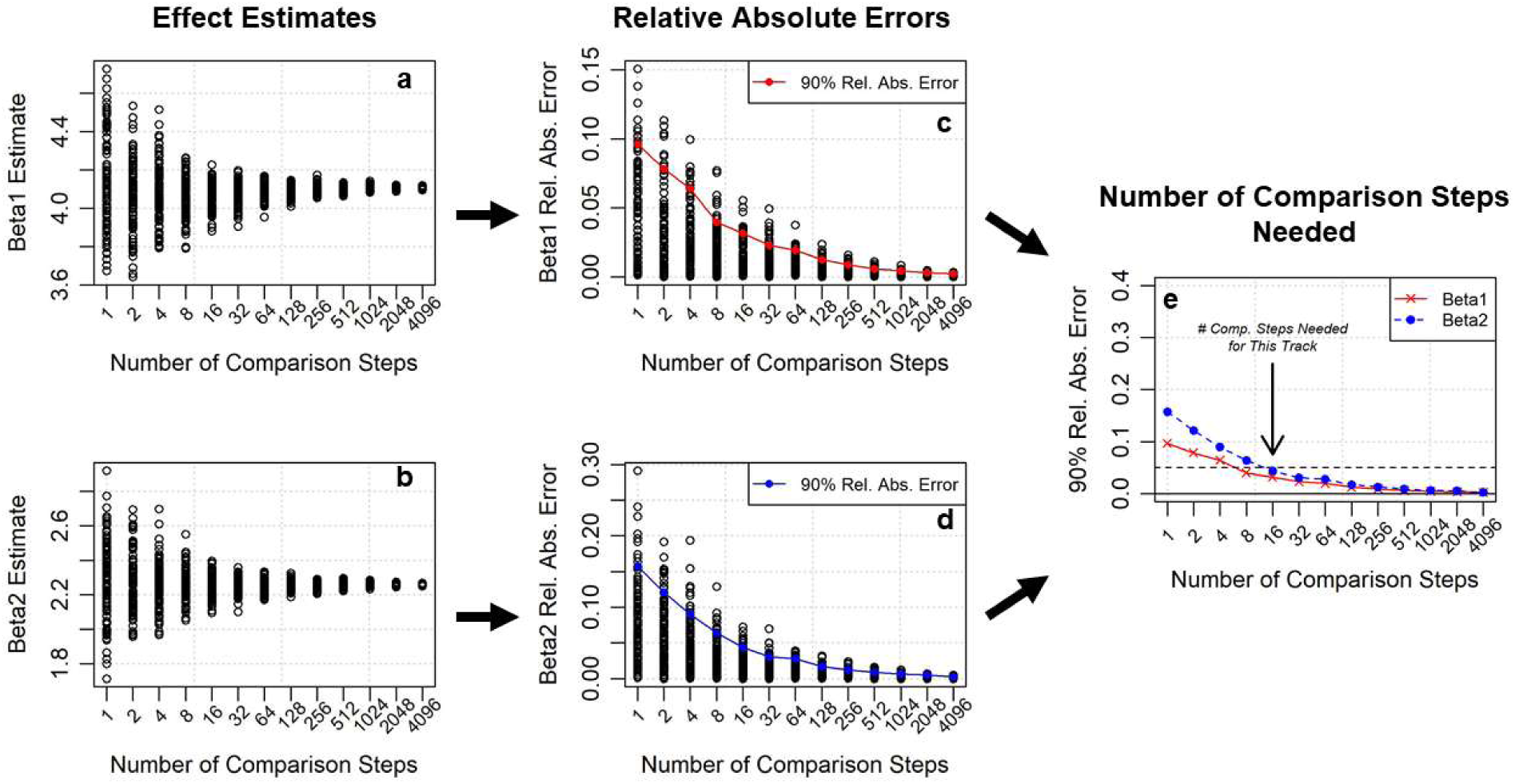
Process to Determine Comparison Steps Needed for a Track: (a,b) estimates of covariate effects, (c, d) absolute errors of estimated effects relative to asymptotic estimates, (e) analysis of comparison steps needed per observed step for this track. Note that 8 comparison steps are needed for the 90% relative absolute error of covariate 1 to fall below 0.05, whereas 16 comparison steps are needed for both covariates 1 and 2 to meet that threshold. All horizontal axes are on a logarithmic scale to more easily distinguish values with small numbers of comparison steps.

From the *q^i^_2,M_* values, we compute *q_2,M_* = max*q^i^_2,M_* (see Fig. 2E) and find the minimum *M* for which *q*_2,M_ ≤ s_1_ for track *j* (across all covariates *i*), denoting this number of CSs as *M_j_\**. With *M_j_\** CSs per OS, at least 100*(1-*ε_2_*) = 100*0.9 = 90 models will have |*RE*_i_|≤ *ε_1_*. When this inequality is met, we regard the track’s SSF as having “converged” and *M_j_\** as the number of CSs “needed” for this track *j*.

We also look for trends among tracks in the same class (see Table 1). We compute the 100*(1-*ε_3_*)^th^ quantile of the *M_j_\** values over all 25 tracks, denoting this quantile as *M\**. For this study, we also set *ε_3_* = 0.1. For a given class of tracks, this amounts to having no more than 2 tracks with *M_j_\** > *M*.

Taken together, these settings allow the identification of the value of *M* that affords consistent accuracy in the estimation of SSF effects. With *ε_1_* = 0.05 and *ε_2_* = *ε_3_* = 0.1, we identify the number of CSs for which 90% of tracks have 90^th^ quantiles of RAEs less than 0.05 (for all covariates involved). For a given covariate, Fig. 2 illustrates the progression from an individual track’s estimated effects and relative errors to the number of CSs needed for a given track. We have selected values of *ε_1_*, *ε_2_*, and *ε_3_* that correspond to common thresholds. Other researchers, however, may select different values that better reflect their research priorities.

To enable more granular comparisons between track classes, we also compute the mean 90^th^ percentile RAE, *A*_Q_ = mean(*q_2,M_*), across all values of *M* and all tracks in a given class. *A*_Q_ is akin to an “area under the curve” obtained by plotting log_2_(*M*) vs. *q_2,M_*. Comparing such areas under the curve is a common practice for comparing model results; see Walter et al. (2015) for an example. While *M\** provides practical insights for the number of CSs needed for a particular track or class of tracks, it is also tied to the specified values of *ε_1_*, *ε_2_*, and *ε_3_*. On the other hand, *A*_Q_ more generally considers the effect of *M* on RAEs of estimated effects and enables more detailed comparison of the influence of landscape and track characteristics on effect estimation.

## 3. Results

Within our simulated tracks, it is clear that the number of CSs influences the variance and potentially also the bias of effect estimates (see Fig. 3). With fewer CSs, estimated effects vary by 35% or more of the magnitude of the effect. Depending on the particular random sample of small numbers of CSs paired with each OS, the estimated effect may be close to the true effect or may have considerable bias. With greater numbers of CSs, variance decreases and the particular random sample of CSs becomes less important, leading to more reliable estimates of covariate effects.

**Fig. 3.**
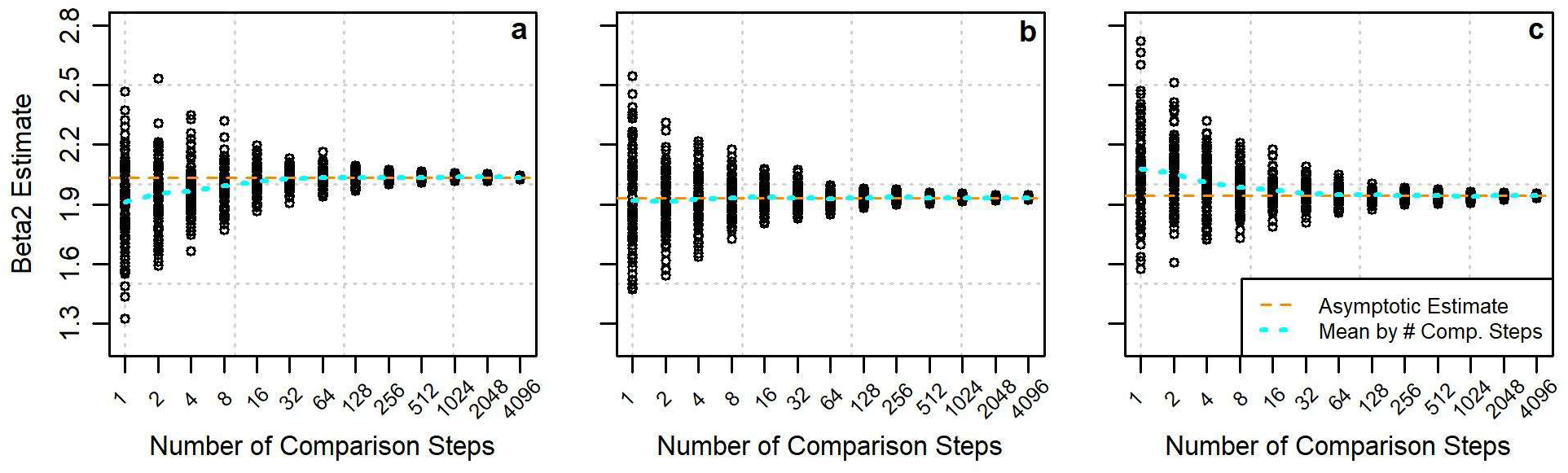
Examples of Effect Estimates by Number of Comparison Steps with Initial (a) Negative Bias, (b) No Bias, (c) Positive Bias

Using the approach described above, we computed the number of CSs needed for each simulated track. We selected a value that allows 90% of replicates to have a RAE less than 5%. **Error! Reference source not found.** shows an example of the number of CSs needed for each covariate for a single simulated track within our base class of tracks. In this figure, the curve for *β*_1_ falls below the 5% line with 8 CSs per OS, while the curve for *β*_2_ falls below the 5% line with 16 CSs. Thus, 16 CSs per OS are needed for this track. The plots for the covariates for other base-class simulated tracks are in the Supplemental Material (Appendix F).

While the track shown in **Error! Reference source not found.** requires 16 CSs per OS to consistently achieve RAE less than 5%, other tracks within the base class require 32 CSs. Fig. 4 shows the distribution of CSs needed for the 25 simulated tracks within this class. For achieving RAEs less than 5% with such tracks, 32 CSs per OS suffice. Based on this result, we suggest *M* = 32 as a reasonable number of CSs if a more detailed analysis of the optimal value of *M* is not feasible.

**Fig. 4.**
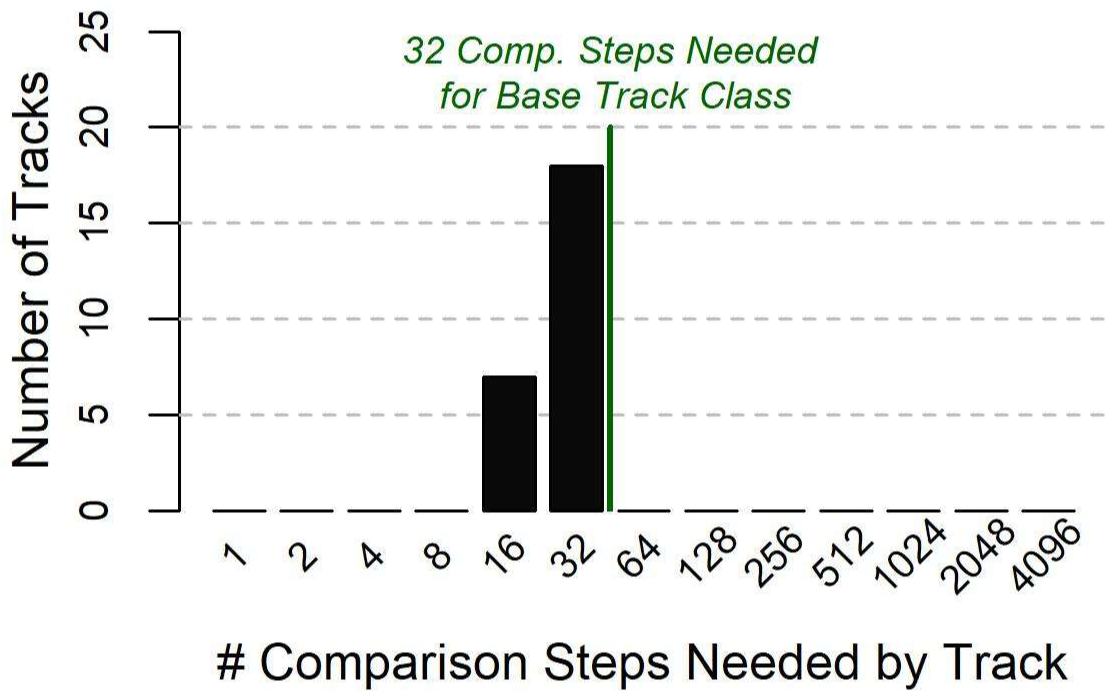
Distribution of Comparison Steps Needed by Track for Base Class of Tracks

We repeated this analysis for classes of tracks with different landscape smoothing window sizes, specified mean step lengths, or track lengths. Table 2 summarizes the influence of these changes upon the number of CSs needed, revealing distinct trends. These results suggest that tracks with relatively greater landscape smoothing, shorter mean step lengths, or shorter track lengths require more CSs to achieve the same level of accuracy in estimates of covariate effects. Longer tracks require fewer CSs. These results can similarly inform researchers decisions about what value of *M* is appropriate for their SSF study. See the Supplemental Material (Appendix F) for plots of distributions of CSs needed for tracks within different classes.

**Table 2.**
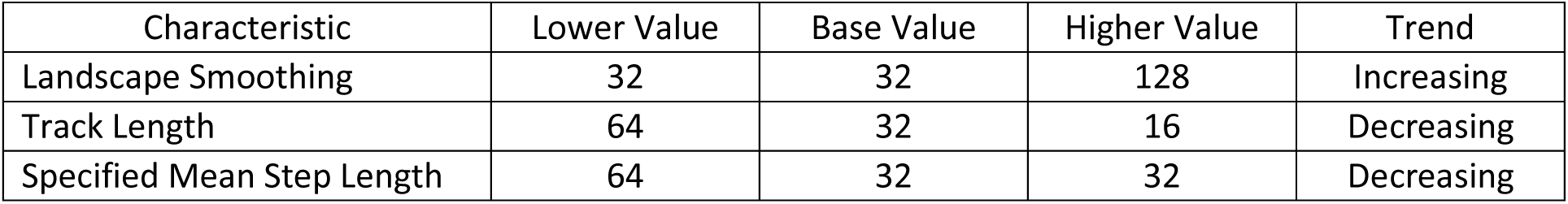
Number of Comparison Steps for Each Class of Tracks.

| Characteristic | Lower Value | Base Value | Higher Value | Trend |
| --- | --- | --- | --- | --- |
| Landscape Smoothing | 32 | 32 | 128 | Increasing |
| Track Length | 64 | 32 | 16 | Decreasing |
| Specified Mean Step Length | 64 | 32 | 32 | Decreasing |

To examine these trends with greater resolution, we also computed the mean 90^th^ percentile RAE across all numbers of CSs and all tracks within a given class. These values do not directly assess how many CSs are needed for individual tracks, but they enable even comparison of the influence of landscape and track characteristics on effect estimation accuracy. These values confirm the trends observed in Table 2. Greater effect estimation errors arise for tracks with relatively greater landscape smoothing, shorter mean step lengths, or shorter track lengths. See Table S-5 in the Supplemental Material for detailed results.

This analysis informs how many CSs are needed for a particular track, potentially enabling large savings in computational requirements and runtime. For instance, Fig. 5 shows the mean runtime per replicate for each track within our base class of tracks (which includes 1000 OSs) with different numbers of CSs per OS. For a single replicate, the mean runtime with our recommended 32 CSs per OS was 8.6 seconds; with 4096 CSs per OS, the mean runtime was over 8 minutes. For a study with 30 individual animals (assuming only one replicate per track), this reduction in runtime amounts to nearly four hours on average. Given the possible need to repeat and troubleshoot analysis, this savings can be quite valuable.

**Fig. 5.**
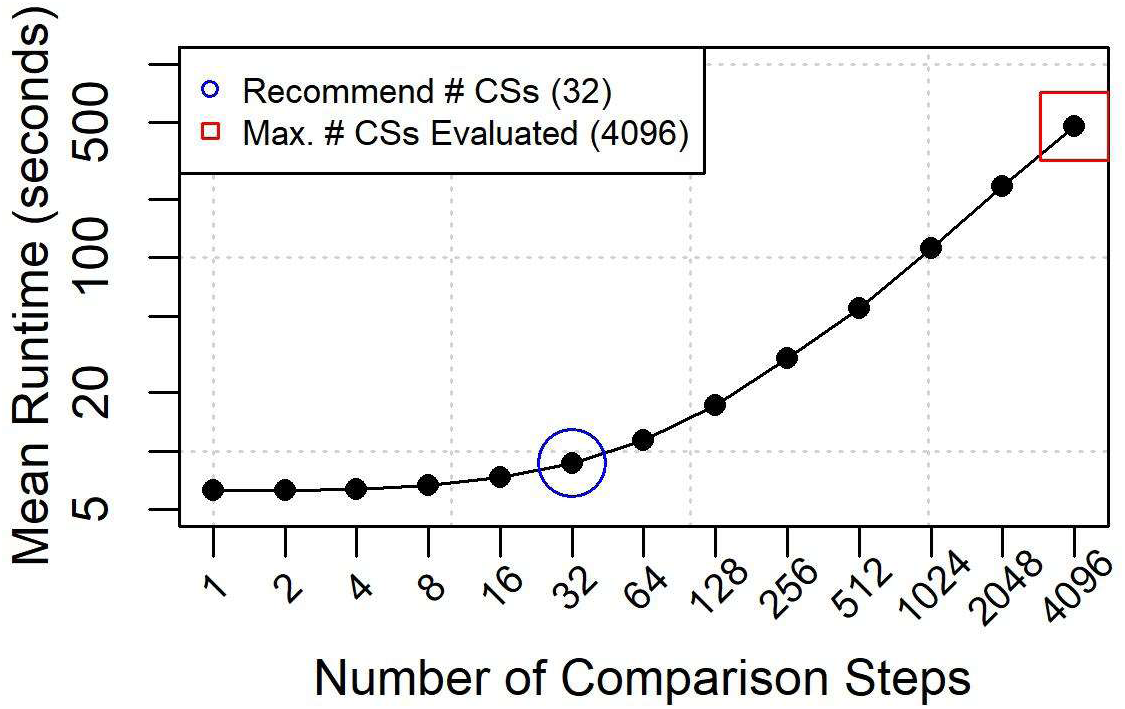
Impact of Number of Comparison Steps per Observed Step on Step Selection Function Runtime (plotted with both axes on a logarithmic scale)

## 4. Discussion

With this research, we have proposed a rigorous process by which the number of CSs needed for a given animal movement track can be identified. We repeatedly sampled sets of different numbers of CSs, fitting an SSF for each replicate and monitoring how the error in estimated covariate effect changes as the number of CSs increases. Given assumptions about tolerable levels and rates of error, this process identifies numbers of CSs needed for specific tracks and for entire classes of tracks and landscape contexts with specific characteristics.

We applied this process to determine the number of CSs needed for simulated tracks with particular characteristics. For tracks in our base class of tracks (which featured modest step lengths and track durations in moderately smooth landscapes), we determined that 32 CSs would limit RAE in covariate estimates to 5% or less for at least 90% of replicates within 90% of tracks and landscapes with these characteristics. For tracks and landscapes with varied characteristics, we found that between 16 – 128 CSs would enable this level of accuracy in estimating covariate effects. In contrast to some existing recommendations and current practice, we did not find any support for either very small or very large numbers of CSs.

Our conclusions regarding numbers of CSs needed for our simulations are based on selected parameters that specify tolerable error levels (*ε_1_*) and frequencies with which these levels are exceeded for individual tracks (*ε_2_*) and classes of tracks (*ε_3_*). We sought to select values that correspond to a high level of accuracy and consistency, while still accounting for discrepancies arising from particular tracks and sets of CSs. Depending on researcher priorities, however, alternative values of *ε_1_*, *ε_2_*, and *ε_3_* could be specified.

Within our analysis, we used *ε_1_* = 0.05. To check the influence of this selection, we also conducted our analysis with *ε_1_* = 0.01, keeping *ε_2_* = *ε_3_* = 0.1. As expected, this analysis determined that larger numbers of CSs are needed to consistently achieve lower error levels. For the base class of tracks, 1024 CSs for each OS are needed to achieve 1% relative error levels with 90% of replicates on 90% of all tracks. This large number of CSs needed illustrates how setting the error threshold *ε_1_* to a lower value generally leads to an increase in CSs needed. See the Supplemental Material (Appendix F) for additional details on these results.

Additionally, we focused on the RAEs of effect estimates to assess when a number of CSs suffices. An alternative metric would be the relative standard deviation (RSD) of the estimates within all models fitted with a given number of CSs. For a given covariate *z*_i_ with the standard deviation of its estimates as *σ*_i_, we compute *RSD_i_* = σ_*i*_/*β^*^_i_*. (This value is similar to a coefficient of variation but differs in that it involves dividing by the simulated effect β^*^_*i*_ rather than the mean of the estimates.) For a fixed number of CSs, this metric quantifies the extent to which an effect estimate will vary based on how the particular CSs are sampled. In the Supplemental Material (Appendix F), we present results for which we require the RSD to fall below a specified threshold, rather than RAE. When using RSD as the convergence criterion, the number of CSs needed is slightly smaller than but comparable to the corresponding number of CSs when using RAE.

This study has several limitations stemming from the need to limit our scope to a reasonable number of combinations. We did not assess the influence of number of covariates, magnitude of specified effects (*β_1_* = 4 and *β_2_* = 2 for us), assumed turning angle distribution (von Mises) and parameter (*κ_A_* = 3), assumed step length distribution (exponential), or autocorrelation between steps (potentially handled with GEE). Likewise, we only evaluated a small number of mean step lengths (*μ_L_*, = 2, 4, or 8), landscape smoothing window sizes (*W* = 1, 2, or 4), and track lengths (*N* = 500, 1000, or 2000). Varying any of these settings could provide an avenue for subsequent research that would meaningfully assess the robustness of our findings. These limitations should be kept in mind when applying our findings for future SSFs.

Moreover, our research shares limitations that are inherent with any SSF. Within our simulations, we do not specify a particular fix frequency or spatial scale, instead assuming that time between fixes reasonably corresponds to the time required for an animal to traverse (i.e., be displaced by) the simulated step lengths. In practice, though, fix frequency may substantially influence the success of SSFs. If fixes are too infrequent, legitimate steps may be missed. On the other hand, if fixes are more frequent, consecutive fixes may not reflect actual steps and may not involve any actual movement decision. As high-frequency tracking data are becoming increasingly available, Munden et al. (2021) propose a novel method for down-sampling fixes at irregular intervals in order to best reflect when an animal’s course has turned, i.e., when an animal has made a movement decision. Ultimately, further work is needed to assess the interaction between fix frequency and the number of CSs needed for an SSF.

While these results do not encompass all possible tracks traversing all landscapes, they provide a starting point for any researcher seeking to determine a reasonable number of CSs for constructing an SSF. For instance, we did not find any classes of tracks for which fewer than 16 CSs sufficed, nor any individual tracks for which fewer than 8 CSs were needed. Regardless of landscape and track characteristics, we found that reliably accurate effect estimation requires at least 16 or 32 CSs. Thus, when fitting SSFs with fewer than 16 CSs per OS, researchers should recognize that their estimates of covariate effects may not be reliable.

The process that we have described in this paper is computationally demanding. We sampled CSs for and fitted 1300 models for each track (see Section 2.3), spanning more than 27 hours of runtime on average per track within our base class. However, while we hope that researchers will benefit from the findings related to particular numbers of CSs discussed in the previous paragraph, we do not intend for researchers to regularly follow the process that we used to obtain those numbers. (By the time that an SSF would have been fit with 4096 CSs per OS, researchers could have reasonable confidence that the resulting effect estimates were accurate. The remaining 1299 models would not be necessary.) Instead, we offer the results of this process as a starting point for other researchers. Moreover, the process that we have described enables farther-reaching benefits as a possible source of objective “ground truth” for the number of CSs needed for a given track.

Herein, as we lay out our approach for how we determined how many CSs are needed for particular simulated tracks, we likewise aim to pave the way for a general strategy for determining an appropriate number of CSs. Given an arbitrary animal track, such a strategy would concretely recommend how to determine how many CSs should be paired with each observes step in order to construct an SSF that achieves specified accuracy levels for each covariate’s estimated effect. Further work is needed to assess possible strategies. Such strategies might include iteratively fitting small numbers of SSFs with increasingly many CSs, as with Stratmann et al. (2021). Nevertheless, our efforts show how the ground-truth values for tracks’ sufficient numbers of CSs that will be needed for such an assessment might be determined.

If the number of animals, track length, and covariate space are sufficiently small or the available computational resources are sufficiently large, we can simply pair very many CSs (e.g., 4000 or 5000) with each OS. In such a case, our proposed process of assessing of how many CSs will suffice is not necessary. However, when the numbers of animals, OSs per animal, or covariates pose potential computational demands, this kind of analysis can be of particular value.

## 5. Conclusions

We have demonstrated a process by which the number of CSs needed for a given animal movement track can be determined. Using this process, we found that 32 CSs per OS are needed for simulated tracks in our base class of tracks. For simulated tracks with other characteristics, we determined that between 16 – 128 CSs per OS are needed. These results provide a starting point for other studies that aim to fit SSFs to animal tracks. Moreover, our process provides the foundation for additional studies that may identify strategies that can be tailored to particular tracks. In general, our results clearly illustrate that the number of CSs influences how well SSFs estimate covariate effects and that different numbers of CSs are appropriate for different tracks and landscape contexts. Researchers should take care to select the number of CSs paired with each OS because this number will ultimately affect the accuracy of effects estimated by their models.

## Declarations

### Ethics approval and consent to participate

Not applicable

### Consent for publication

Not applicable

### Availability of data and materials

The dataset supporting the conclusions of this article is available in the following repository: https://github.com/StatsForAnimalMovement/InsightsForSSFs/. Code for analyzing these results is also included as a supplement to this paper.

### Authors’ contributions

PK conceptualized the study and designed its methodology, wrote and validation the simulation and analysis, curated the simulated data, and wrote the manuscript. All authors reviewed and edited the manuscript. WF provided supervision and aided interpretation of the results.

## Acknowledgements

Not applicable

## Authors’ information

PK is a doctoral student in the Applied Statistics track of the Applied Mathematics, Statistics, and Scientific Computing program at the University of Maryland.

## List of Abbreviations

CLR: Conditional logistic regression
CS: Comparison step
GPS: Global positioning system
OS: Observed step
RAE: Relative absolute error
RSD: Relative standard deviation
RSF: Resource selection function
SSF: Step selection function

## Supplemental Material

### Appendix A. More Detail on Step Selection Functions

Step selection functions (SSFs) are underpinned by the following biological hypothesis: Given an animal’s current location, the likelihood that the animal will step to at another particular location is related to the conditions at the other location or more generally along the path between the locations. In particular, SSFs aim to characterize the factors that motivate the animal to “select” a step (which has been observed within the track) rather than any other steps (which have not been observed). By comparing the observed steps (OSs) with other feasible “comparison steps” (CSs), SSFs provide a statistical framework quantify the effects of these motivating factors.

#### A.1. Notation

We use this biological hypothesis to provide context for introducing relevant notation. At time *t*, we assume that the animal’s current location is ***x_t_***. For terrestrial animals, ***x_t_*** corresponds to two-dimensional longitude and latitude coordinates. The measured conditions at a given location and a given time are ***R***(***x_t_***), which we simplify as ***R_t_***. Often, these conditions involve resources such as vegetation levels or land cover type, but they may also include other factors such as weather, terrain, presence of predators, or proximity to human infrastructure. We measure *V* separate conditions at each location and ***R_t_*** = (*R*_1,t_, …, *R*_V,t_) with *R*_v,t_ = *R*_v_(***x_t_***). The likelihood that the animal will be next observed at location ***x_t_***_+**1**_ is related to the conditions at the destination of, along, or surrounding the path from ***x_t_*** to ***x_t_***_+**1**_. For a given step, we compute covariates ***z_t_***_+**1**_ for each step that concludes at ***x_t_***_+**1**_, i.e., that has ***x_t_***_+**1**_ as its destination. We use similar notation for covariates as for conditions with ***z_t_***_+**1**_ = ***z***(***x_t_***, ***x_t_***_+**1**_) = (*z*_1_(***x_t_***, ***x_t_***_+**1**_), …, *z*_P_(***x_t_***, ***x_t_***_+**1**_)) = (*z*_1,t+1_, …, *z*_P,t+1_). The number of measured conditions *V* does not necessarily equal the number of computed covariates *P*.

For this analysis, we consider only the conditions at the destination of a step; i.e., for each *p*, *z*_p,t+1_ = *R*_v,t+1_ for some *v*. Given this convention, we refer to each step simply by its destination. For example, we refer to the step with ***x_t_***_+**1**_ as its destination simply as ***x_t_***_+**1**_. We discuss additional options for computing covariates below. In this analysis, we also assume that all conditions are static at a given location and do not change in time. Relaxing this assumption increases the computational demand of the analysis but generally does not affect the approach.

In attempt to infer the effects of factors that motivate the animal’s movement decisions, we compare the characteristics of OSs with the characteristics of unobserved but feasible CSs. Each OS is paired with *M* CSs. We use ***x^m^_t+1_*** to denote the *m*th CS paired with step ***x_t_***_+**1**_. The covariates associated with this step are ***Z^m^_t+1_*** = (*Z^m^_1,t+1_*, …, *Z^m^_P,t+1_*). The set of the ***x^1^_t+1_***, …, ***x^M^_t+1_*** paired with a given OS ***x_t_***_+**1**_ comprise a stratum; this stratum also includes ***x_t_***_+**1**_. Recommending a number of CSs within each stratum is the focus of this analysis. In summary, *t* indexes the OSs and strata, while *m* indexes the CSs within a given stratum. For consistent notation, we refer to the OS within each stratum also as ***x^0^_t+1_***.

The CSs originating from a given location ***x_t_*** are drawn from a distribution *F* of feasible alternative steps. This distribution is typically expressed in terms of step length (displacement) and relative turning angle (change in bearing). This distribution may directly consist of observed pairs of step lengths and relative turning angles from other animals in the study. In other cases, these data from other animals may be fitted to a parametric distribution (Thurfjell et al. 2014). Thus, the distribution of alternative steps may be generally denoted as *F*(γ) with parameters γ. In Fig. 1 from the main paper (reproduced as Fig. S-1), the CSs that have been paired with the animal’s third OS (shown in blue) are shown as red dashed arrows.

**Fig. S-1.**
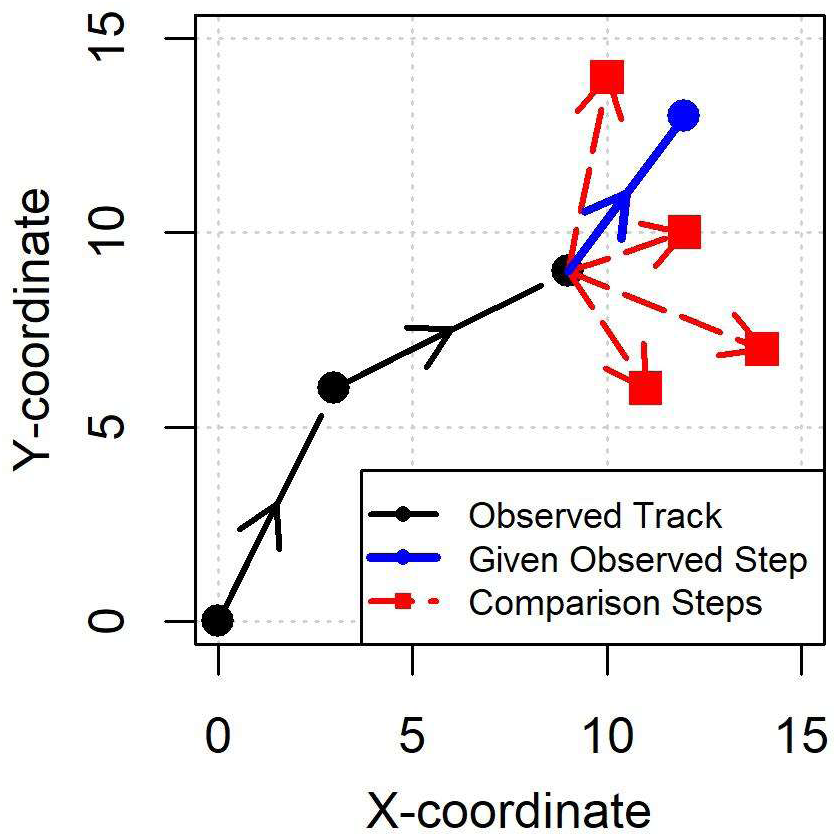
Steps within an Example Animal Track

#### A.2. Motivation

This analysis strategy is based on two assumptions. First, we assume that an animal has some internal “function” that consistently guides its decisions about which next “step” to select. This function incorporates all of the factors or “slices” of the nearby environment that affect the animal’s movement decisions, as well as the animal’s past experiences. Secondly, we assume that we can track enough of the environmental covariates and the animal’s movement history in order to infer this function. Fig. S-2 illustrates these assumptions.

**Fig. S-2.**
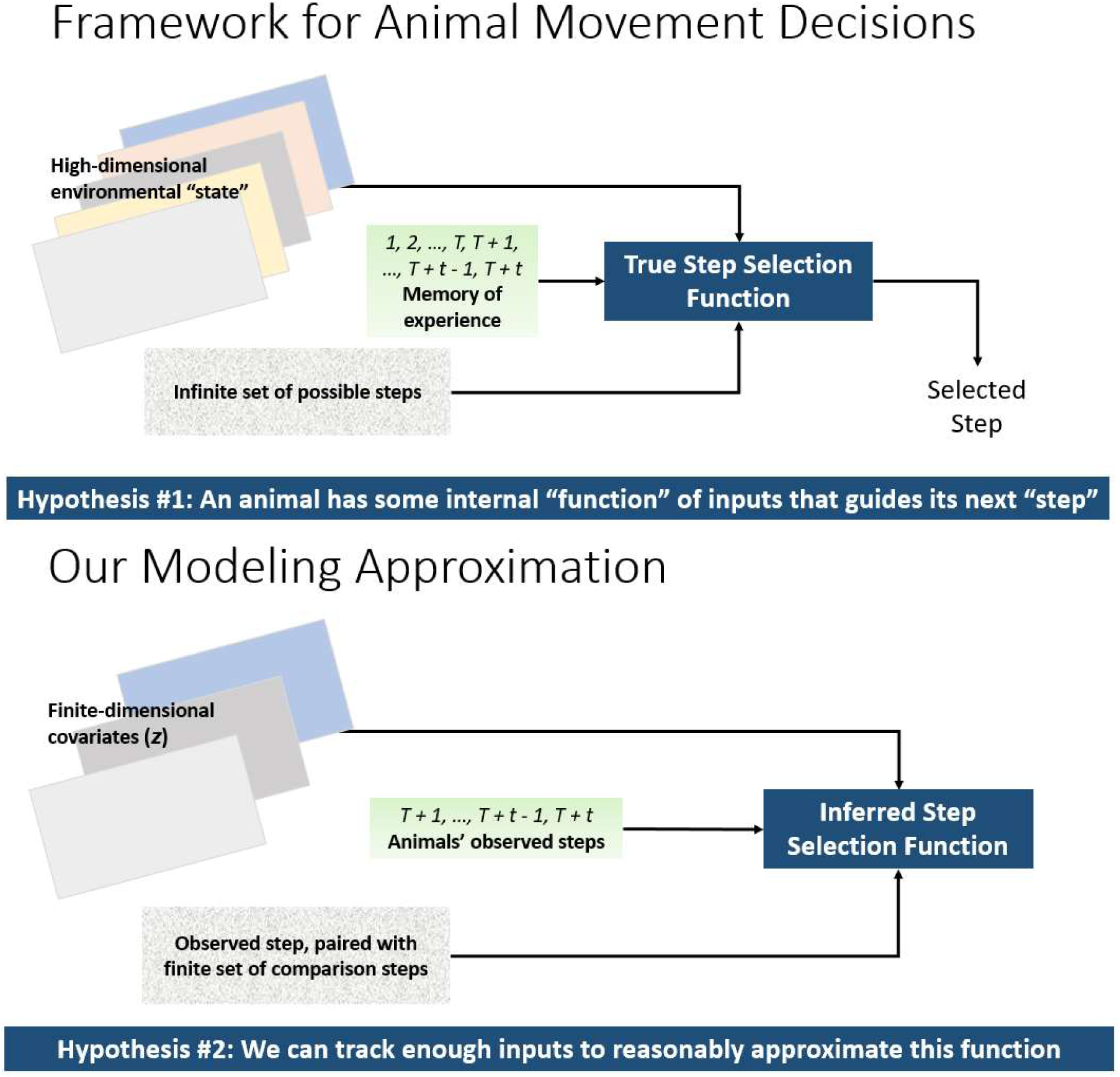
Step Selection Function Framework

#### A.3. Modeling Decisions

SSFs typically have an exponential form, conditioned upon the current location and previous step ***x_t_***:

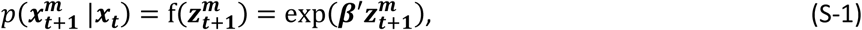

where ***β*** are coefficients (“effects”) fitted to the covariates in order to model whether a given step has been observed.

These effects reflect how strongly each resource or environmental factor or step characteristic influences the movement of a particular animal or set of animals. Such results are often interpreted in light of previous hypotheses regarding the movements of particular groups of animals. For instance, Fortin et al. (2005) used SSFs to illustrate how shelter (aspen stands) and predator presence (wolves) affect elk movement.

Previous researchers have measured environmental covariates (e.g., terrain, land cover, weather, predator presence) in different ways. In some cases, the covariates have been measured at the end of a given linear step (i.e., at the tip of the arrows in Fig. S-1):

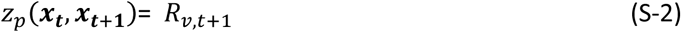

Alternatively, covariates may be calculated from multiple points along the straight line connecting the two endpoints of a step (i.e., spanning the entire length of the arrows in Fig. S-1):

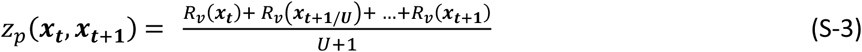

In this equation, *U* is a constant integer reflecting the measured factor’s spatial resolution. For any value δ within [0,1], we use ***x_t_***_+ẟ_ = (1 − δ)***x_t_*** + δ***x_t_***_+**1**_ to indicate a point in between ***x_t_*** and ***x_t_***_+**1**_. Some studies, e.g., Avgar et al (2016), have calculated covariates both at endpoints and along the linear step. As another approach, the assumption of linear steps can be relaxed by considering distributions of potentially nonlinear paths between starting and end points. Kriging estimates such distributions (Fleming et al. 2016).

Moreover, in this analysis, we have assumed that resources and other conditions relevant to step selection are static. In practice, these values may vary in time. Nevertheless, the approaches for computing covariates described in this section can also be applied to time-varying data.

#### A.4. Different Objectives for Step Selection Functions

Different researchers use SSFs with different objectives. Most but not all SSF studies focus on identifying or characterizing factors that affect animal movement; other studies have used SSFs to predict space use. Relevant factors are identified through several methods including model selection and effect significance (e.g., determined by comparing each covariate’s p-value with a significance threshold). Criteria for model selection include Akaike information criterion (AIC) (Akaike 1973), corrected AIC (AICc) (Sugiura 1978; Hurvich and Tsai 1989; Burnham and Anderson 2022), and Watanabe-Akaike information criterion (WAIC) (Watanabe 2009). Table S-1 summarizes different ways in which researchers have leveraged SSFs. Several studies have used AIC or another criterion to select a best model, then compared the effects of the covariates within that model.

**Table S-1:**
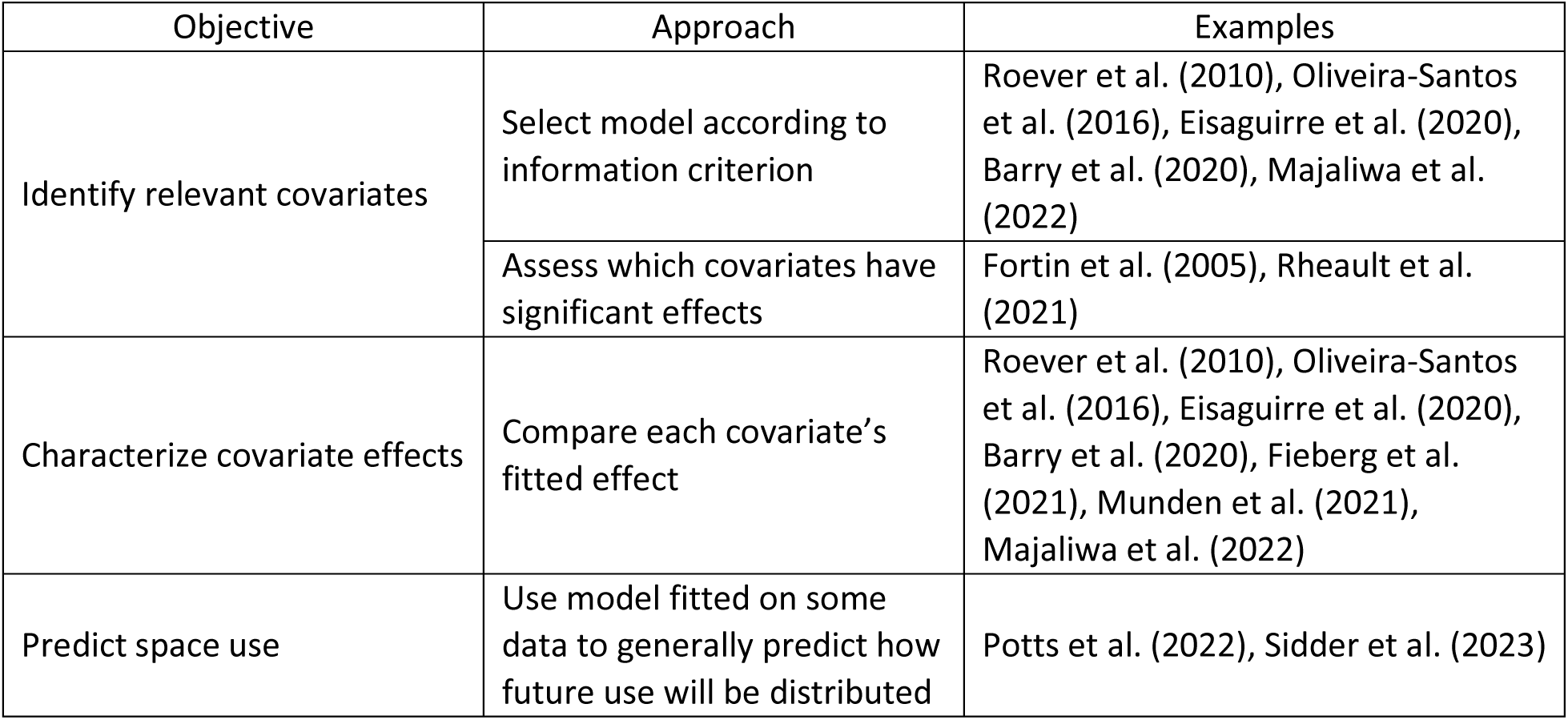
Possible Objectives of Step Selection Functions.

| Objective | Approach | Examples |
| --- | --- | --- |
| Identify relevant covariates | Select model according to information criterion | Roever et al. (2010), Oliveira-Santos et al. (2016), Eisaguirre et al. (2020), Barry et al. (2020), Majaliwa et al. (2022) |
|  | Assess which covariates have significant effects | Fortin et al. (2005), Rheault et al. (2021) |
| Characterize covariate effects | Compare each covariate's fitted effect | Roever et al. (2010), Oliveira-Santos et al. (2016), Eisaguirre et al. (2020), Barry et al. (2020), Fieberg et al. (2021), Munden et al. (2021), Majaliwa et al. (2022) |
| Predict space use | Use model fitted on some data to generally predict how future use will be distributed | Potts et al. (2022), Sidder et al. (2023) |

### Appendix B. Model Selection with Additional Comparison Steps

Depending on the particular objective for an SSF study, different metrics are appropriate to determine how many CSs are needed. In the main paper, we focused on the objective of characterizing covariate effects. Accordingly, we use relative absolute error (RAE) and relative standard deviation (RSD) of effect estimates to assess how many CSs are needed for a given track. In this section, we instead focus on the objective of identifying relevant covariates through model selection.

Typically, a “best model” is selected by comparing the values of a criterion such as AIC for a set of candidate models. The AIC of each fitted model with *P* parameters is computed directly from the model’s log-likelihood *LL*.

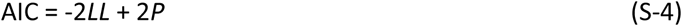

Because of the additional degrees of freedom, models with additional covariates tend to have larger log-likelihood values (yielding smaller values of -2*LL*) but are penalized for the additional covariates.

Given three covariates, there are eight possible subsets of covariates. In a preliminary analysis (not described here), we did not observe any models with subsets of covariates that yielded the lowest AIC other than the two “true” covariates or the “full” set of all three covariates (with those two covariates and a third “noise” covariate). Within our study, we assess whether the correct model with the two true covariates has a lower AIC than a model with all three covariates. We refer to these values of AIC as *AIC2* and *AIC3* respectively. If *AIC2* < *AIC3*, i.e., if *AIC2* – *AIC3* < 0, the correct model is selected. On the other hand, if the model that includes the noise covariate has the lower AIC, that model will be incorrectly selected.

In general, we anticipate that models fitted with more CSs per OS will have lower incorrect selection rates. If desired, we could assess how many CSs are needed in order to have satisfactorily low incorrect selection rates (e.g., below 5% or 1%). However, as described below, we found that models fitted with more CSs per OS do not always yield lower incorrect selection rates. This phenomenon greatly complicates the use of incorrect selection rates for determining the CSs needed for a given track.

Fig. S-3 provides an example of *AIC2* – *AIC3* values for a given track. Each data point corresponds to a model fitted to a separate set of CSs. Points below the dashed line (corresponding to *AIC2* – *AIC3* = 0) indicate that the correct model will be selected. To more easily distinguish the values for models from small numbers of CSs, the horizontal axis is shown with logarithmic scale.

**Fig. S-3.**
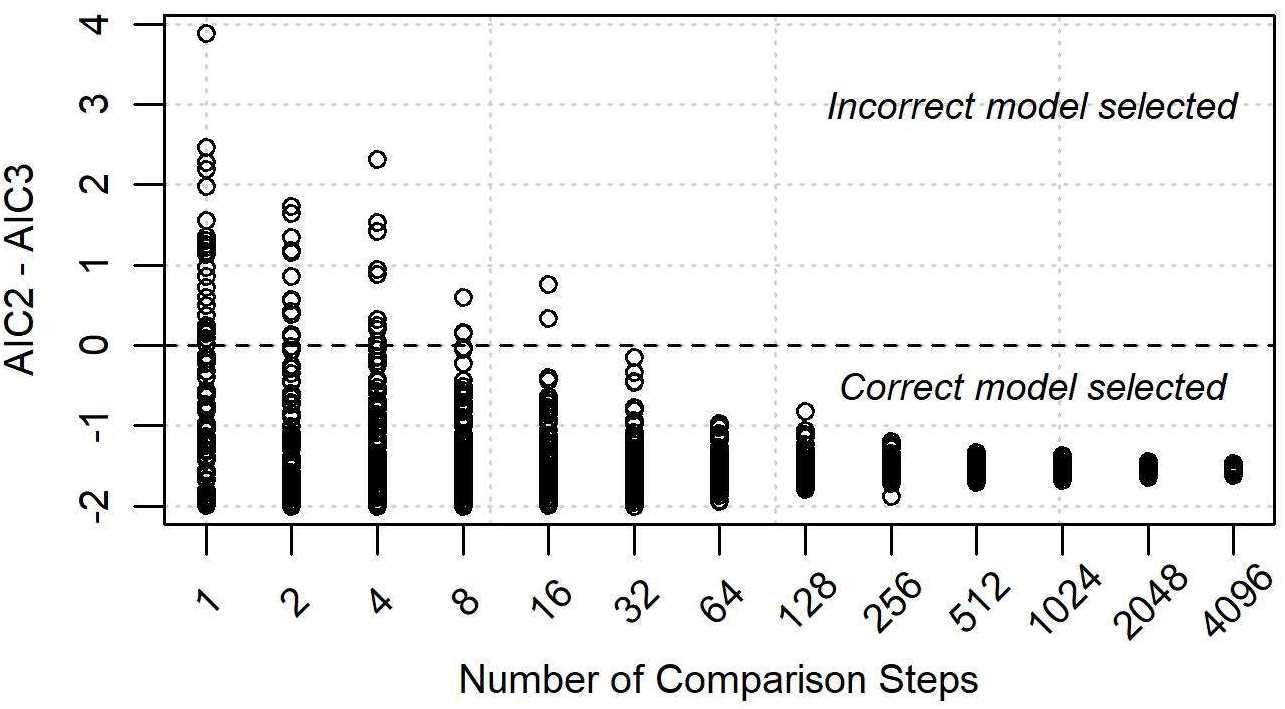
AIC Comparison for Each Replicate for an Example Track

For each number of CSs, we computed the rate of incorrect model selection, e.g., the proportion of models with values above the dashed line in Fig. S-3. These proportions (corresponding to the same example) are shown in Fig. S-4. As expected, incorrect selections for this model decrease as the number of CSs increase. The number of CSs needed for this track depends on the threshold to which we want to limit the incorrect selection rate. For a threshold of 5% (shown by the dashed line), we need 8 CSs.

**Fig. S-4.**
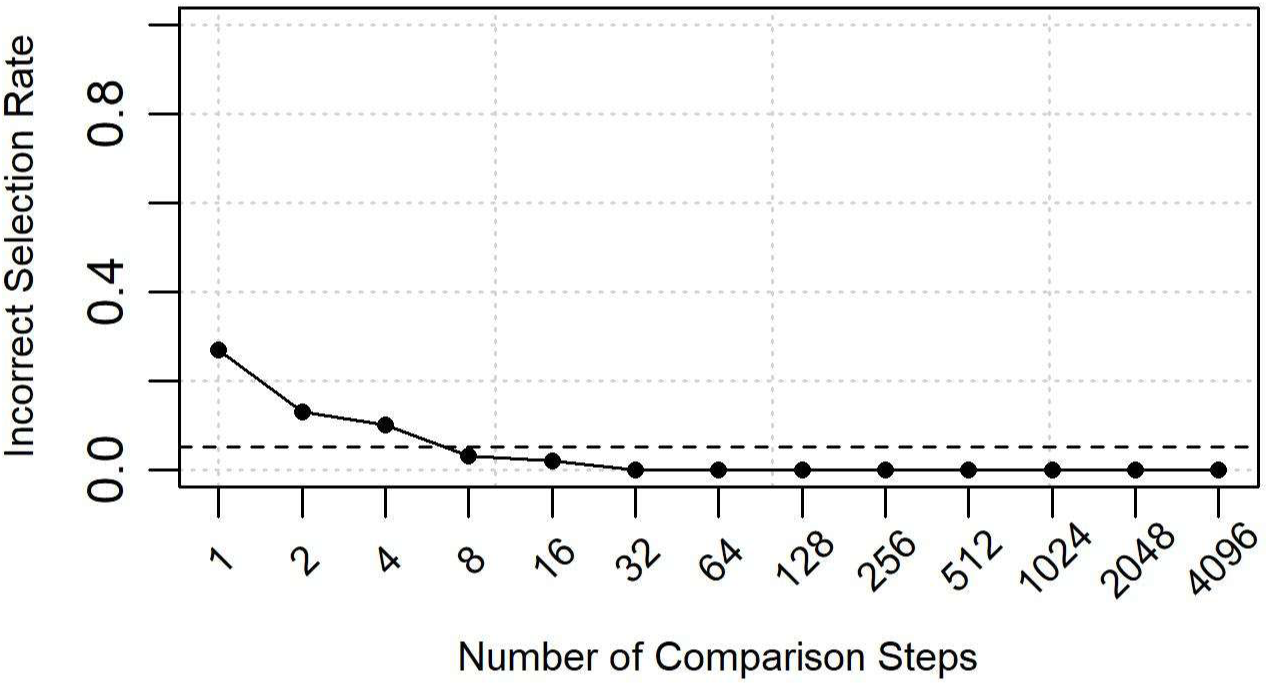
Incorrect Selection Rate for an Example Track, Decreasing as Expected

Most simulated tracks have decreasing incorrect selection rate plots similar to Fig. S-4. However, some simulated tracks have plots of incorrect selection rates with the opposite trend. Unexpectedly, for these tracks, as the number of CSs increases, the rate at which the incorrect model was selected also increases. Fig. S-5 provides an example of such an increasing trend.

**Fig. S-5.**
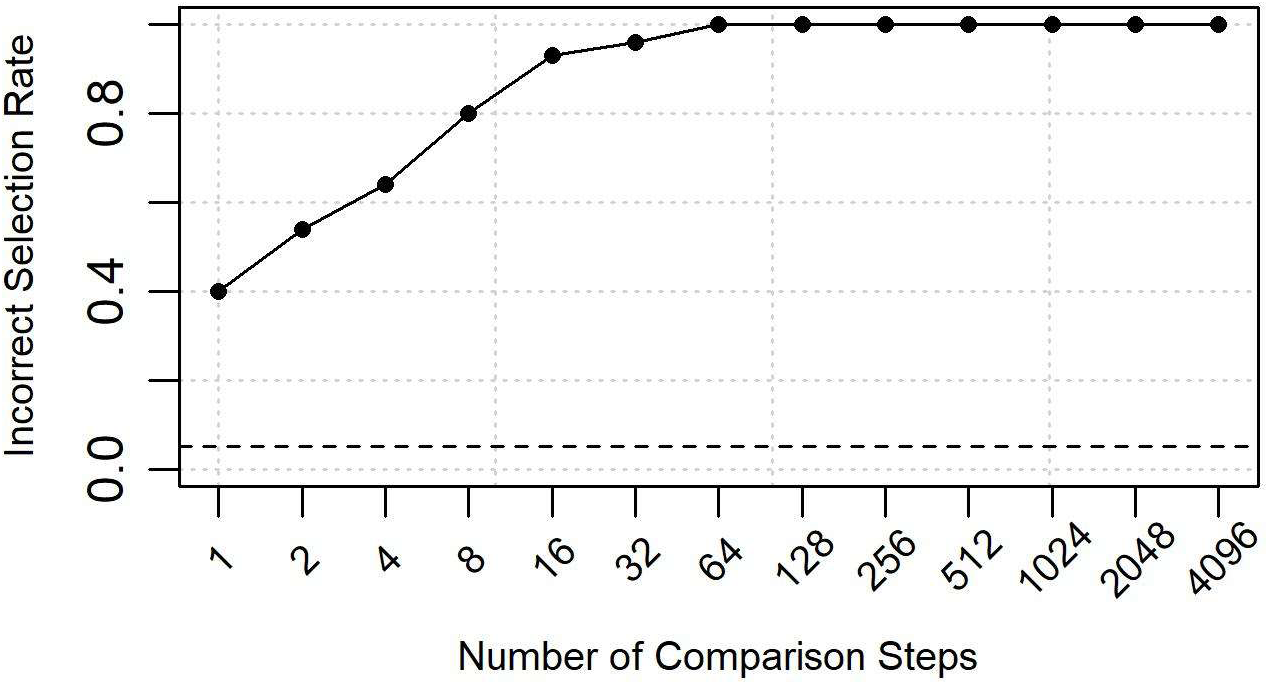
Incorrect Selection Rate for an Example Track, Unexpectedly Increasing

There is no immediately evident difference between these tracks. Both tracks were simulated on the same covariate landscape. As shown in Fig. S-6, both tracks span approximately the same amount of space.

**Fig. S-6.**
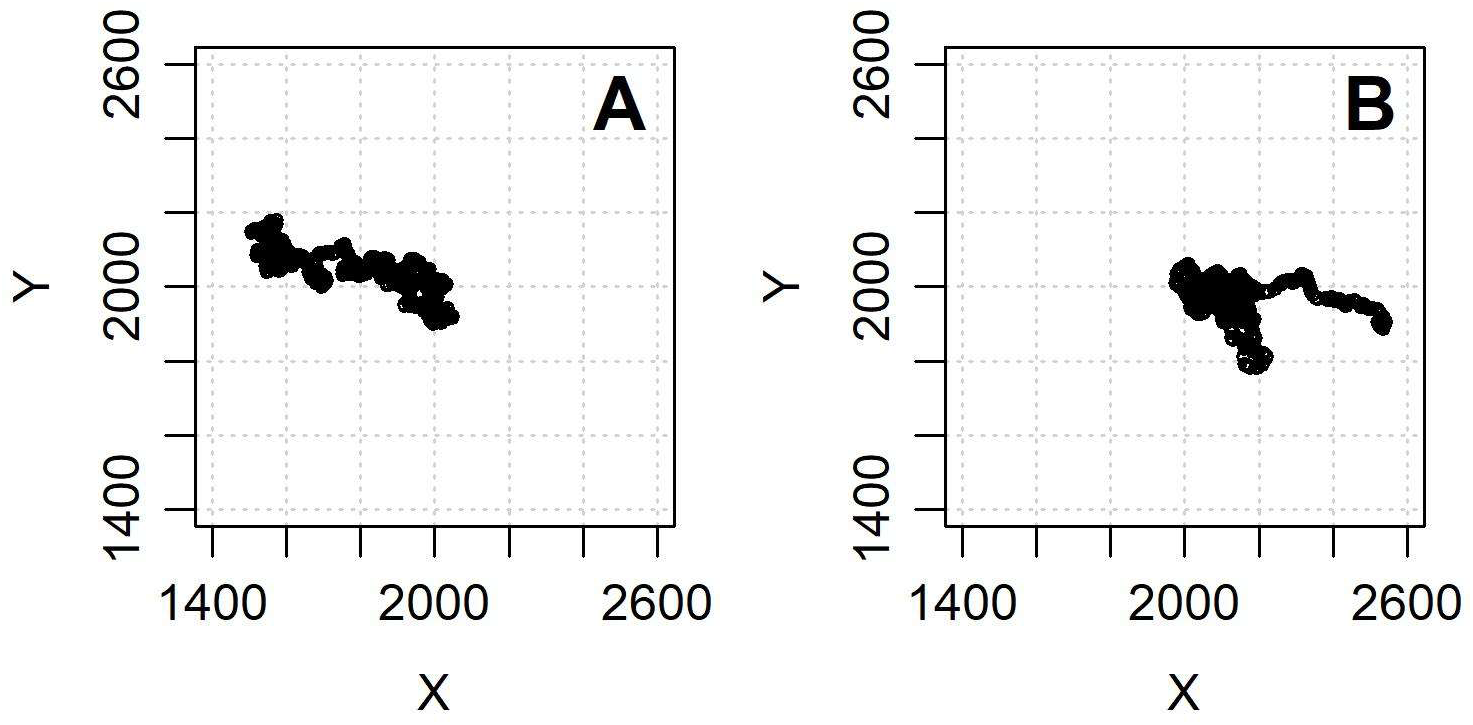
Simulated Tracks Corresponding to the (A) Decreasing and (B) Increasing Trends in Incorrect Model Selection

Increasing trends in incorrect model selection occur in similar proportions across the different scenarios, as shown in Table S-2 and Table S-3. Within this table, tracks are classified according to the incorrect selection rate with models fitted to the largest number of CSs (*M* = 4096), i.e., on the right end of the plot. Tracks with final values below 0.05 have “increasing trends,” while tracks with final values above 0.95 have “decreasing trends.” All other tracks have “mixed trends.”

**Table S-2:** Convergence Trends for Different Track Classes.

| Track Class | # Tracks with ... |  |  | Proportion of Tracks with Non-Decreasing Trends |
| --- | --- | --- | --- | --- |
|  | Decreasing Trends | Mixed Trends | Increasing Trends |  |
| Base | 18 | 2 | 5 | 0.28 |
| Less Smooth Landscape | 20 | 0 | 5 | 0.20 |
| More Smooth Landscape | 22 | 0 | 3 | 0.12 |
| Shorter Track | 20 | 1 | 4 | 0.20 |
| Longer Track | 17 | 1 | 7 | 0.32 |
| Shorter Steps | 20 | 2 | 3 | 0.20 |
| Longer Steps | 21 | 0 | 4 | 0.16 |

**Table S-3:** Proportions of Tracks with Non-Decreasing Trends by Characteristic.

| Characteristic | Low Value | Base Value | High Value | Trend by Characteristic |
| --- | --- | --- | --- | --- |
| Landscape Smoothing | 0.20 | 0.28 | 0.12 | None |
| Track Length | 0.20 | 0.28 | 0.32 | Increasing |
| Mean Step Length | 0.20 | 0.28 | 0.16 | None |

As described above, there can be two tracks simulated on the same landscape for which one track has a decreasing trend in incorrect selection rate and the other track has an increasing trend. This suggests that the phenomenon stems from the track rather than the landscape. Accordingly, we also considered other characteristics of tracks that might contribute to the increasing trend. These characteristics include: simulated step lengths (possibly differing from the specified step length); correlation between the noise covariate and one of the covariates with a specified true effect; variation in noise covariate values along the simulated track; and spatial extent spanned by the simulated track.

Fig. S-7 shows results of these comparisons across all tracks from all seven track classes (amounting to 175 total tracks). Among the track characteristics, *AbsZ* is the absolute value of the Z-score of the noise covariate with the track. We computed the Z-score as the mean divided by the standard deviation. Since the noise covariate does not influence track simulation, we expect that this Z-score would be close to 0, but with some tracks the absolute value differs from 0. *Cor1* and *Cor2* measure the correlation between the first and noise covariates and the second and noise covariates respectively, while *MaxCor* is the maximum for each track of these two correlation values. *ACF* is the autocorrelation among the time series of values for the noise covariates at successive locations along a given track (i.e., with lag = 1). *MSL* is the mean simulated step length within the track (not necessarily equal to the specified mean step length, see Appendix E). *XRange* and *YRange* are the differences between the minimum and maximum values in the X- and Y-dimensions respectively, while *Area* is their product (essentially the area of a bounding box around the track). We computed correlation between each of these characteristics with a binary indicator *(HighIMS*) of whether the incorrect model selection rate exceeds 5% with the maximum number of CSs (*M* = 4096).

**Fig. S-7.**
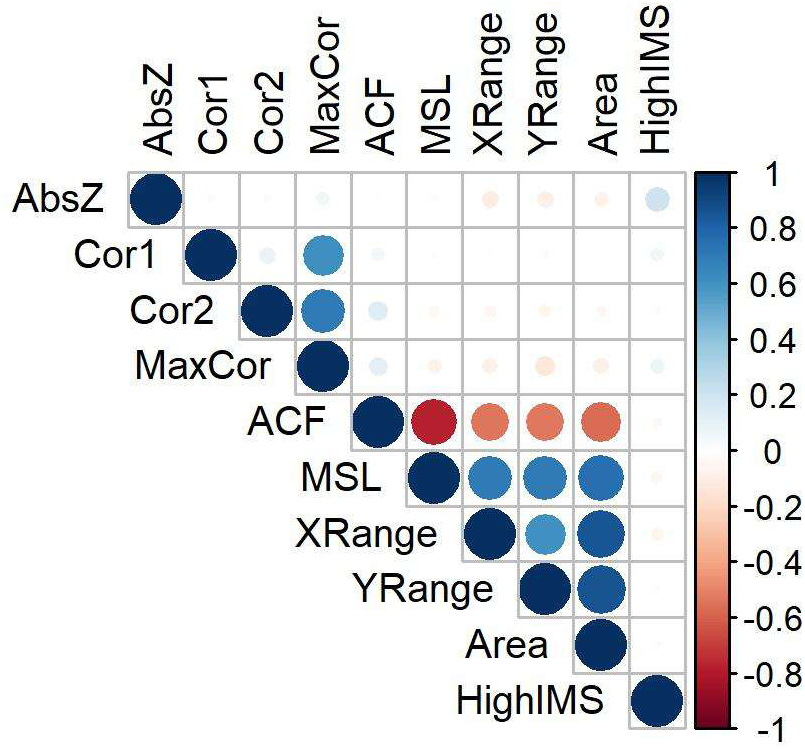
Correlation Between Track Characteristics and Increasing Trends in Model Selection Errors

This analysis fails to identify any clear explanations of phenomena that may lead to increasing trends in false positives or incorrect model selection. As expected, the correlation values (*Cor1*, *Cor2*, *MaxCor*) are correlated with each other. Likewise, the characteristics related to step length (*ACF*, *MSL*, *XRange*, *YRange*, *Area*) tend to be strongly correlated with each other. However, none of these characteristics show strong correlation with the increasing error trend indicator. The only track characteristic that shows some correlation (0.21) with the increasing error trend indicator is *AbsZ*. Tracks that randomly have noise covariate values with a nonzero bias tend to correspond to models in which the noise covariate is incorrectly included in AIC-based model selection.

These observations are consistent with known risks of incorrectly identifying “noise” covariates as relevant via AIC-based model selection. Sutherland et al. (2023) cautioned that otherwise uninformative covariates can appear to be significant or can be included in the selected model due to those covariates having values that are somewhat correlated to the response due to randomness. Given these risks, we encourage researchers to use AIC-based model selection for SSFs only with caution.

We focused on the use of AIC to determine the best model and subsequently identify relevant covariates. In practice, researchers may use other model selection criteria, including AICc or WAIC. Further work is needed to confirm whether our findings extend to model selection involving these alternate criteria.

### Appendix C. Sampling Noise vs. Numerical Error

Animal locations obtained from GPS fixes consist of “presence-only” data for which it is known where an animal has been but not necessarily where it has not been. It is possible that an animal visited a location that was not recorded by a GPS fix. These locations are generally not known to researchers based on the animal’s GPS fixes. Warton and Shepherd (2010) observed that such presence-only data are best modeled with an inhomogeneous Poisson point process (IPPP). Under certain conditions, Fithian and Hastie (2012) showed that the estimates from a logistic regression model converge to the estimates from an IPPP, e.g., with increasingly many CSs or “background samples.”

For each true covariate *z_i_*, we specify an effect *β_i_*. Following the logic of Fithian and Hastie (2012), we assume that our ability to estimate the effect of that covariate within a simulated track is limited when the number of CSs paired with each OS is small but improves as we increase the number of CSs. We refer to the estimation error stemming from insufficiently many CSs as “numerical error” because it is related to efforts to numerically approximate the full set of CSs (i.e., to approximate the denominator in Equation S-5 in Appendix D).

The covariate’s actual effect within a simulated track is also influenced by the randomness in how the track is simulated. In the long run, the covariate’s effect within the simulated track will converge to the specified effect. However, within a track of finite length, the specified effect will be compounded by some amount of noise due to sampling. The actual effect β^*^_*i*_ is the sum of the specified effect *β_i_* and this “sampling noise.” Fig. S-8 illustrates the contributions of both numerical error and sampling noise to the difference between an estimated effect for a given covariate and for that covariate’s specified effect.

**Fig. S-8.**
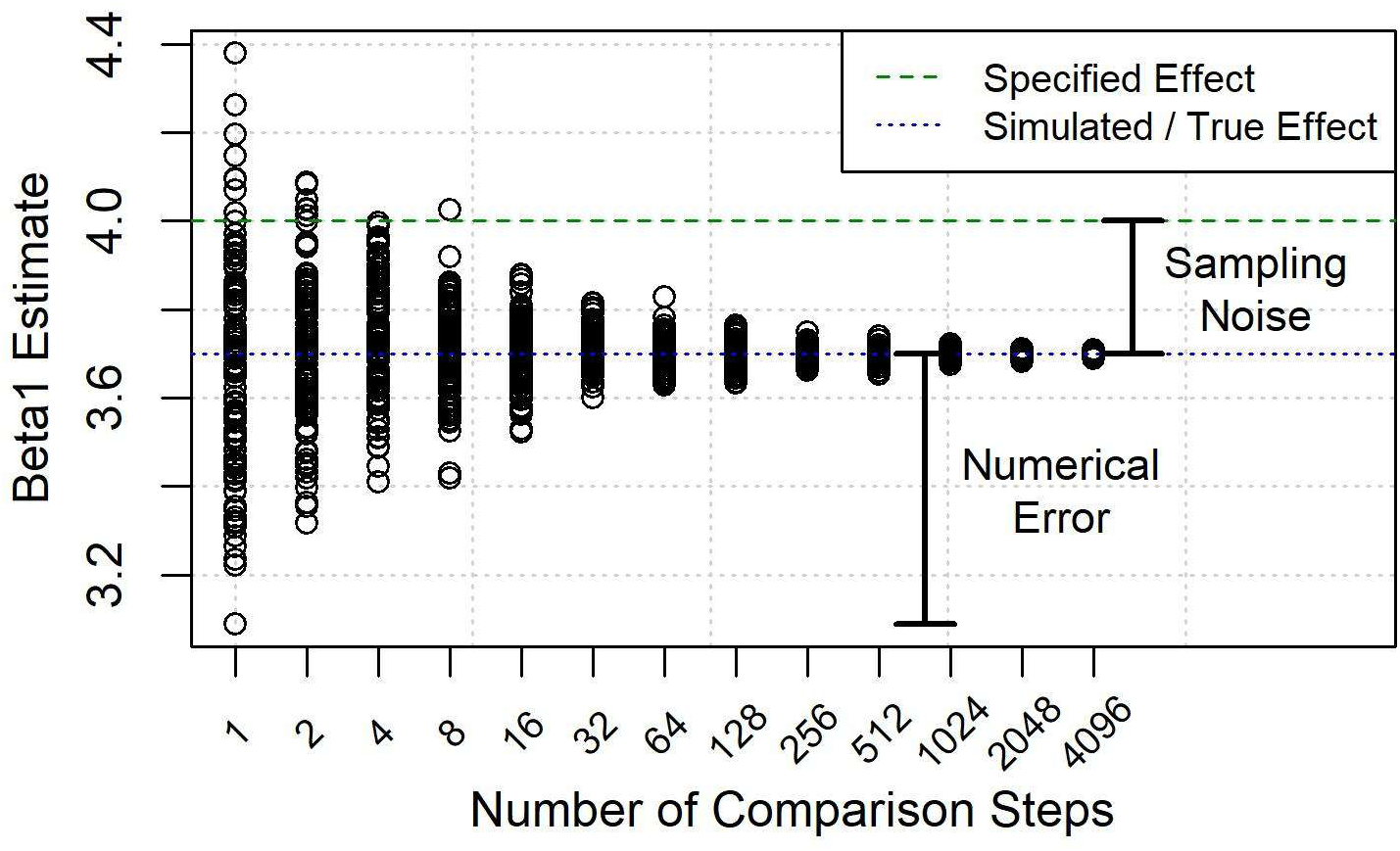
Sampling Noise vs. Numerical Error

Indeed, as track length increases, we observe that sampling noise appears to decrease, i.e., that true effects ***β***^∗^approach specified effects ***β***. Using simulated tracks of lengths 500, 1000, and 2000 steps, we measured the absolute discrepancy between the estimated effect (with 4096 CSs) and the specified effect **β^*^**. As shown in Table S-4, this discrepancy generally decreases as the track length increases. These results come from tracks with specified mean step length *μ_L_* = 4 and smoothing window size *W* = 2, with 25 tracks of each length (five from each of five landscapes) and 100 replicates per track. As in Section 3.2, we specified *β_1_* = 4 and *β_2_* = 2.

**Table S-4:**
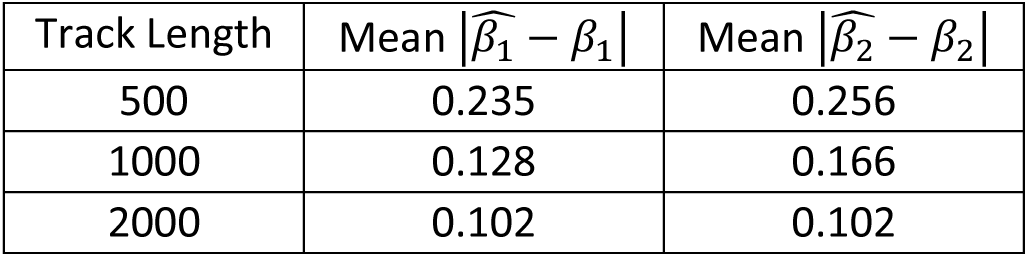
Summary of Sampling Noise with Different Track Lengths.

Additionally, Fig. S-9 shows the estimated mean effect β̂_1_ with 4096 CSs per OS for all tracks with smoothing window size *W* = 2 and mean step length *μ_L_* = 4. Notably, the estimated effects become more concentrated about the specified effect *β*_1_ = 4 as the track length increases.

**Fig. S-9.**
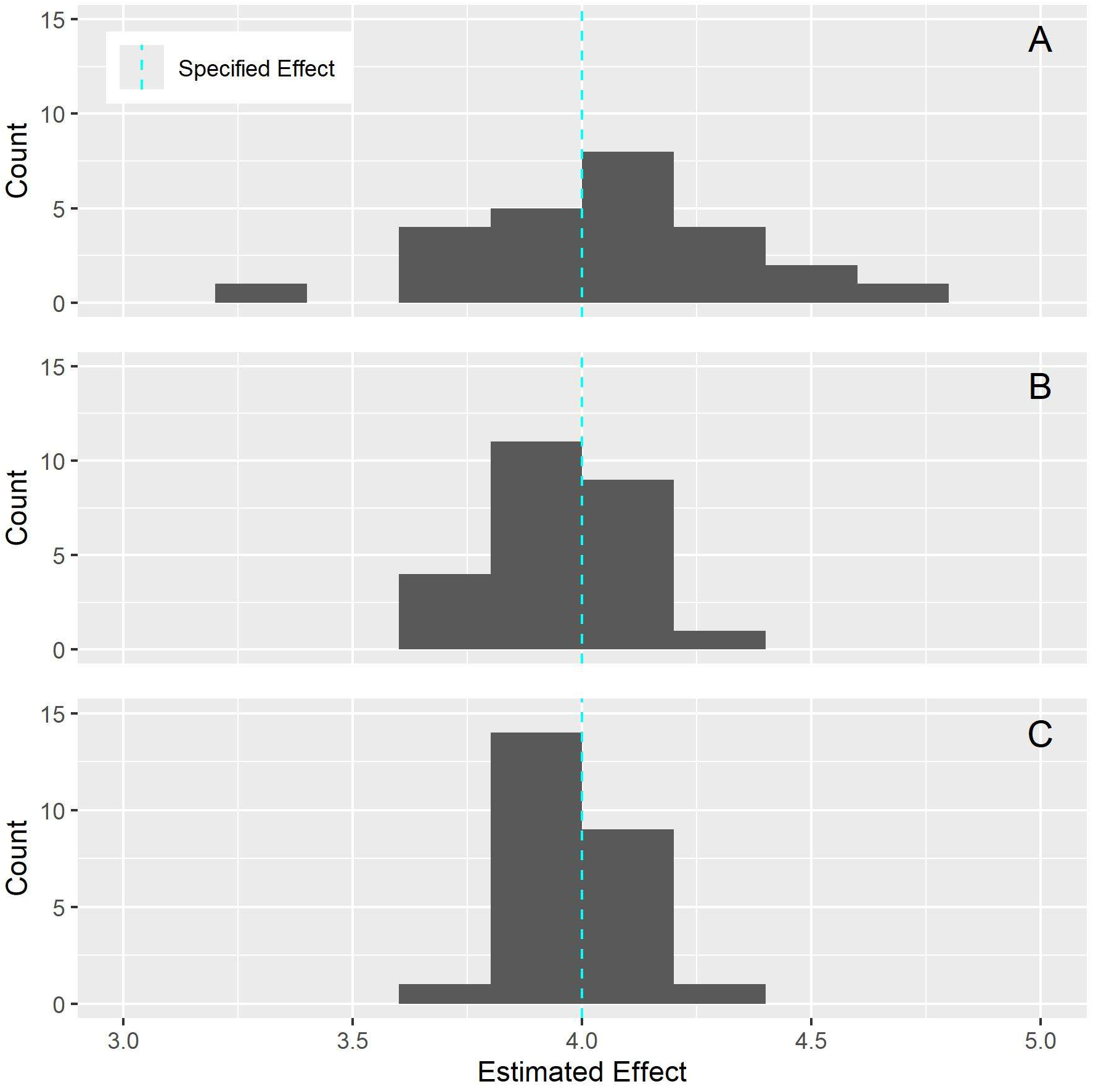
Estimated Effects with Many Comparison Steps for Track Length (A) 500, (B) 1000, (C) 2000

### Appendix D. More Detail on Simulated Data

This section describes our simulation process in greater detail. For completeness, we repeat some detail from the main paper. We first simulate a landscape of covariates, then simulate the tracks within that landscape.

To simulate the landscape of values for a given covariate, we set up a *G*-by-*G* grid of square cells. The grid needs to be large enough to accommodate the full track without the track going beyond the boundary of the grid. Herein, longer tracks may require larger grids. To accommodate our farthest-reaching track, we use *G* = 4001.

For each cell, we sample an initial value from a normal distribution with mean zero and standard deviation one. To achieve different amounts of correlation within the grid, we use a smoothing window with width 2*W*+1. We compute the smoothed value for a given grid cell as the mean of the initial values of the (2*W*+1)^2^ cells within *W* units of the given cell. For instance, when *W* = 1, the smoothed value of one cell is based on the initial values of that cell and eight surrounding cells (i.e., all cells within a three-by-three square). Larger values of *W* achieve greater spatial correlation within the grid. For cells along the boundary of the grid, we consider only the cells within a smoothing window that lie within the grid. The smoothed values for these boundary cells involve the values of less than (2*W*+1)^2^ cells.

Fig. S-10 illustrates this smoothing process, showing the effect of different values of *W*. Currently, we are using *W* = 1, *W* = 2, or *W* = 4. In this example, a 21-by-21 grid of values for a single covariate are simulated. The initial values (panel A) are sampled from a normal distribution with mean zero and standard deviation one. In this figure, all values less than -1 are shown with the same shade of dark blue and all values greater than 1 are shown with the same shade of yellow; such truncation allows differences in intermediate values (which are prominent following smoothing) to be seen in greater detail. Fig. S-11 shows the corresponding histograms of values. With greater smoothing, the histogram follows an increasingly narrow normal distribution.

**Fig. S-10.**
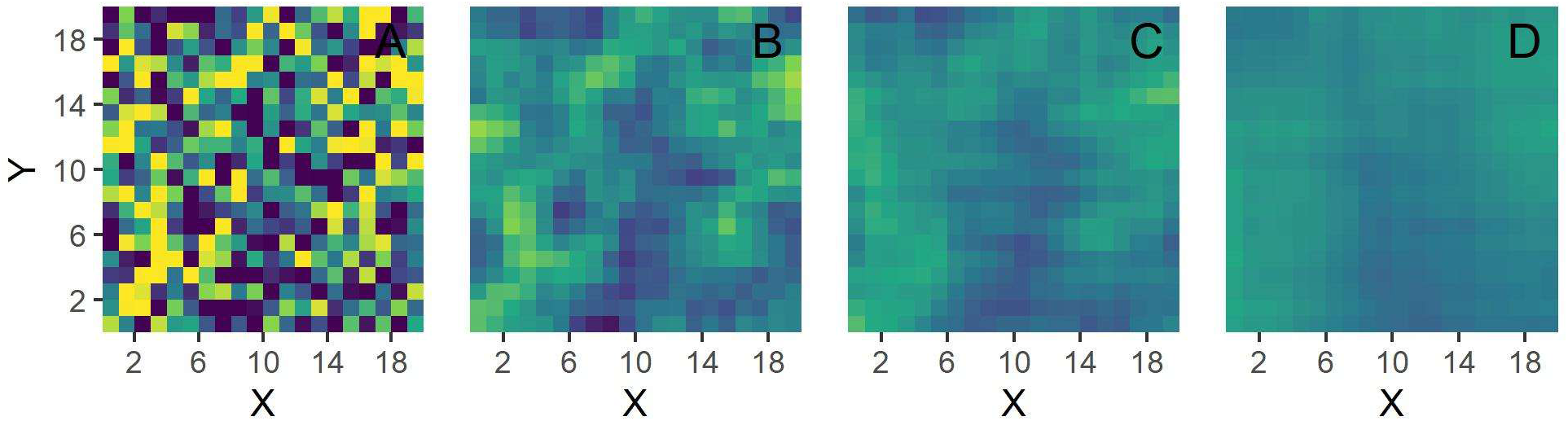
Illustration of Landscape Smoothing, (A) Without Smoothing and With Smoothing Window Sizes (B) W = 1, (C) W = 2, (D) W = 4

**Fig. S-11.**
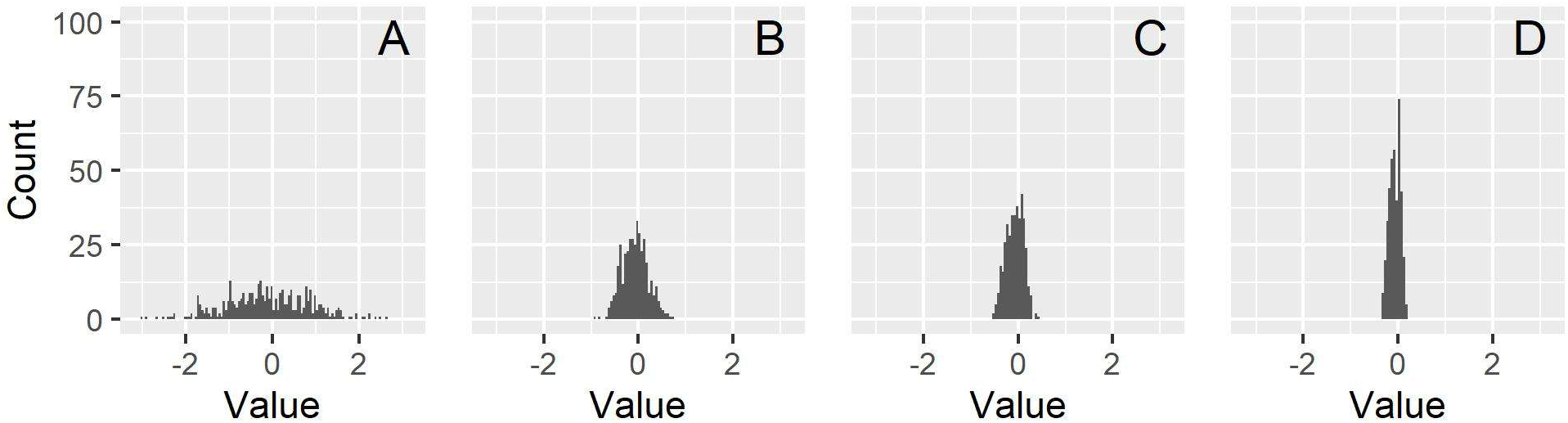
Histogram of Covariate Values, (A) Without Smoothing and With Smoothing Window Sizes (B) W = 1, (C) W = 2, (D) W = 4

We simulate separate grids for each resource (using the value of the resource at a given location as the covariate value for steps that terminate at that location). We do not model any correlation between the grids of different resources. We simulate grids for three resources, using two for simulating tracks and the third to test how the number of CSs affects the capability of conditional logistic regression to select the correct model in the presence of a “noise” covariate (see Appendix B). For studies involving additional simulated covariates, additional grids could be simulated.

Given a simulated landscape, we simulate a track by successively carrying out weighted sampling of candidate steps near the track’s current location; for this study, we consider 10000 candidate steps for each step added to the simulated track. We compute the weights of each candidate step from the covariate values at the step destination. We determine the candidate steps according to randomly sampled step lengths and relative turning angles.

We draw step lengths *L* from an exponential distribution with mean *μ_L_*. We do not specify a unit for step length. Instead, step length can be interpreted according to the spatial resolution of the covariate landscape. For instance, a step length of one corresponds to the width of a single grid within the covariate landscape. We assume that all covariates have the same spatial resolution. Similarly, we draw relative turning angles *A* from a von Mises distribution with concentration parameter *κ_A_*. As default, we use *μ_L_* = 2, 4, or 8 and *κ_A_* = 3 for this study. Correlation between the step length and relative turning angle could be studied for particular animals (e.g., with shorter step lengths corresponding to larger relative turning angles), but we do not include this detail in this study.

We opt to determine covariate values for a given step solely from the resource values at the step’s destination. As a result, the step can be identified by the destination ***x_t+1_*** and the covariate values for that step are simply the values of the environmental factors at the destination, i.e., ***z_t+1_*** = (*z_t+1,1_*, …, *z_t+1,P_*) and for each *p*, *z_t+1,p_* = *R_t+1,v_* for some *v*. An alternative would be to consider the resource values along the entire step, e.g., along the entire line from ***x_t_*** to ***x_t+1_*** (see Appendix A). For each covariate, we specify effects that determine the extent to which that covariate affects that weight of each candidate step. In this study, there are two covariates that affect the track simulation (i.e., *P* = 2); we specify *β_1_* = 4 as the effect of the first covariate and *β_2_* = 2 as the effect of the second covariate. (As described in Appendix C, finite simulated tracks generally do not exactly reflect these specified effects.)

In general, the contributions of these covariates for the *m*th candidate step ***x^m^_t+1_*** are summarized by the resource-based selection probability.

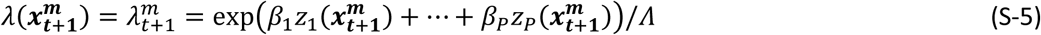

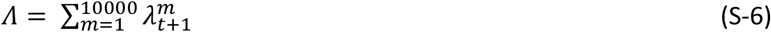

Ultimately, the sampling weights of each candidate step are determined by this selection probability.

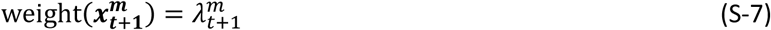

For all CSs that originate from a given OS, we divide their individual weights by the sum of the weights. After this normalizing step, we compute the cumulative weight of the first *m* candidate steps. We uniformly sample a random value between 0 and 1. The candidate step corresponding to the largest cumulative sum less than the random value is the step that is added to the simulated track. (Note that any order of candidate steps can be used with this approach.) In this way, candidate steps with larger weights are more likely but not guaranteed to be sampled. Alternatively, always adding the candidate step with the largest weight leads to issues such as perfect separation for some covariates’ values between OSs and CSs and failure for the model to converge; thus, we use weighted sampling instead.

We continue to iteratively simulate steps according to this process until we have simulated a track with the specified track length *N*+1. The resulting track has *N* OSs, each with *P* covariates. Because a prior step (involving two prior locations) is required to compute relative turning angle, we use *N*-1 OSs for each model. With larger mean step lengths or longer track lengths, tracks cover larger spatial extents. Fig. S-12 and Fig. S-13 illustrate these trends. In Fig. S-12, all tracks have length *N* = 1000. In Fig. S-13, all tracks have mean step length *μ_L_* = 4. In both figures, the tracks have been simulated on landscapes with smoothing window size *W* = 2.

**Fig. S-12.**
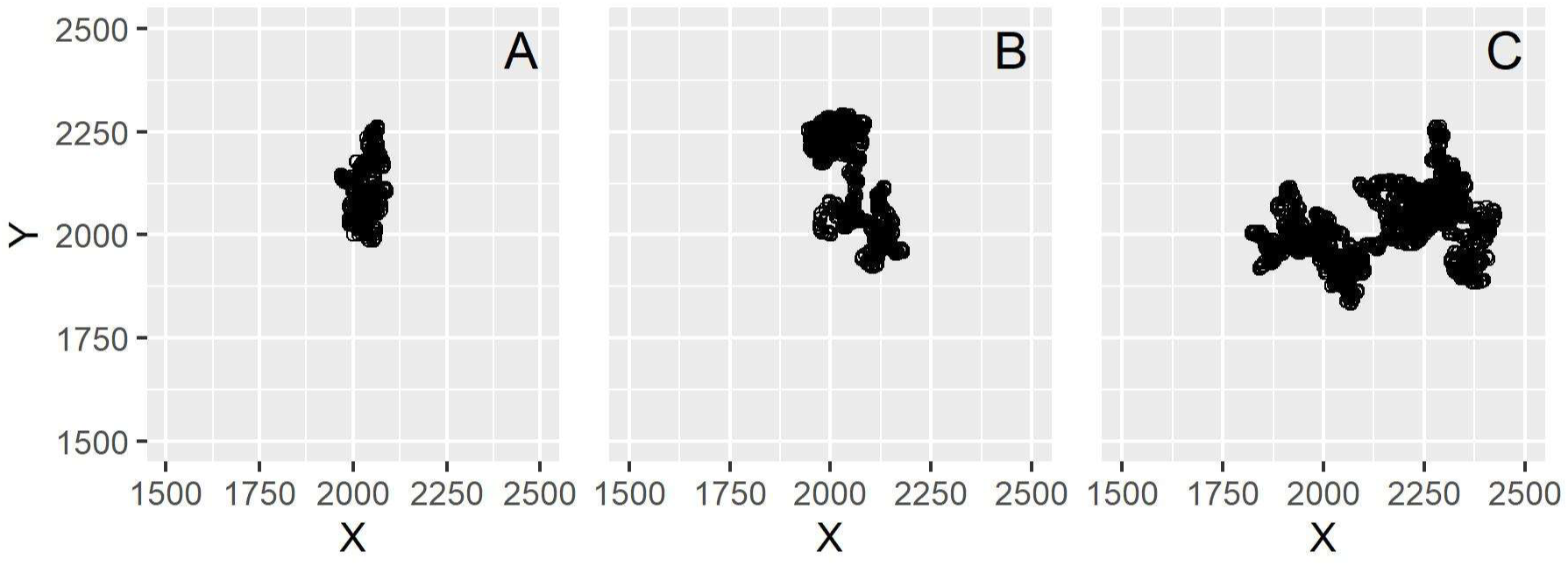
Example Tracks with Increasing Mean Step Lengths, (A) *μ_L_* = 2, (B) *μ_L_* = 4, (C) *μ_L_* = 8

**Fig. S-13.**
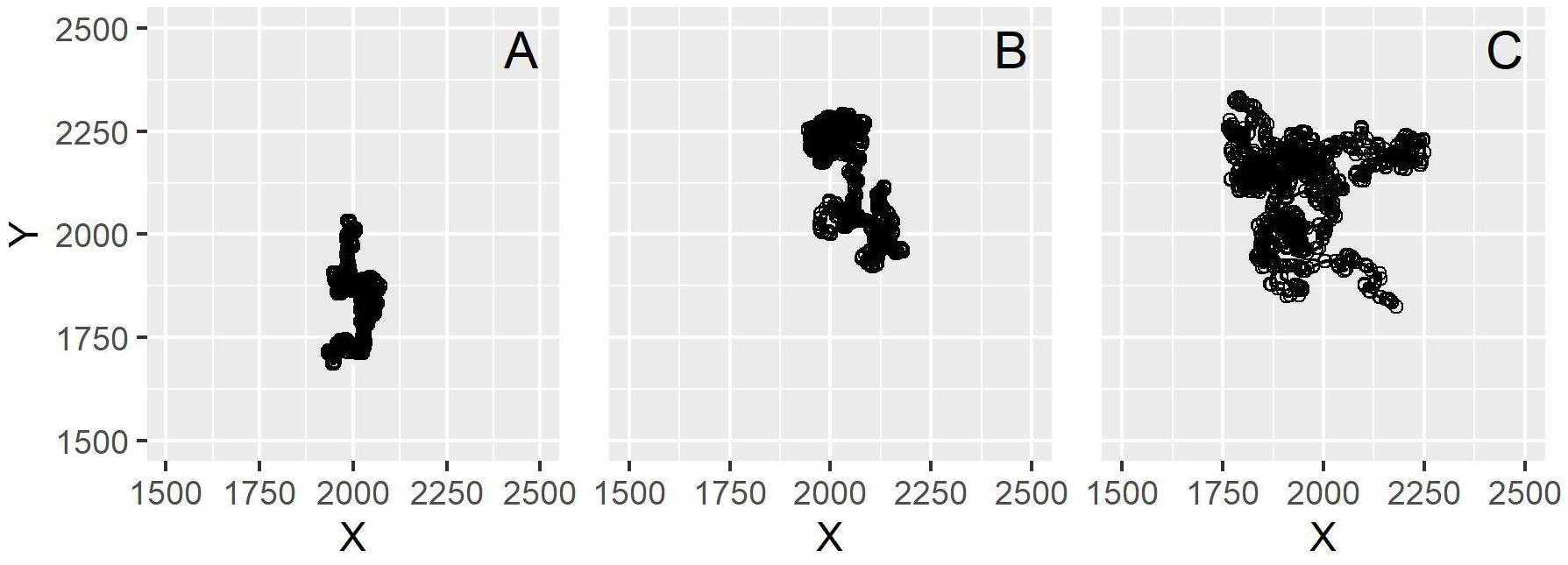
Example Tracks with Increasing Track Lengths, (A) *N* = 500, (B) *N* = 1000, (C) *N* = 2000

### Appendix E. Specified vs. Actual Step Lengths

Within our simulations, we specify a mean step length (*μ_L_* = 2, 4, or 8), but the distribution step lengths that have been actually simulated may differ due to the effect of resource preferences. Because of landscape smoothing (described above), locations with preferable covariate values tend to be near other locations with preferable covariate values. As a result, the actually simulated step lengths tend to be less than what we specified and the distribution of actually simulated step lengths is biased towards smaller lengths. Fig. S-14 shows a histogram of the observed mean step lengths across all 175 tracks within our analysis (spanning the seven track classes described in the main paper). To allow comparison across tracks with different specified mean step lengths, all observed step lengths have been divided by the specified mean step length.

**Fig. S-14.**
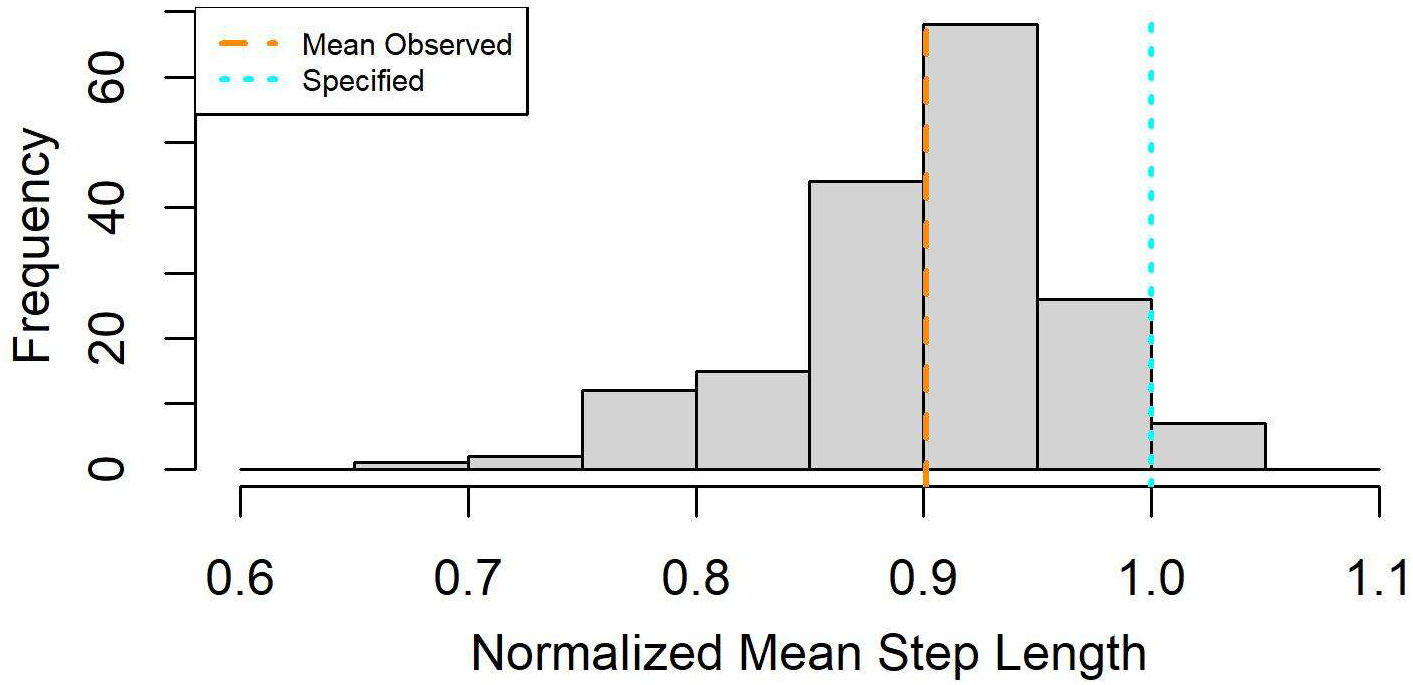
Normalized Mean Step Length by Track

This phenomenon is magnified with less smoothing, e.g., *W* = 1, perhaps due to the increased frequency of tracks remaining in the same cell with preferable covariate values. With greater smoothing, animals can remain near preferable covariate values while taking slightly longer steps. Fig. S-15 compares histograms of normalized mean step length with different smoothing window sizes. Each plot includes a single track class with 25 tracks; each has specified mean step length *μ* = 4 and track length *N* = 1000.

**Fig. S-15.**
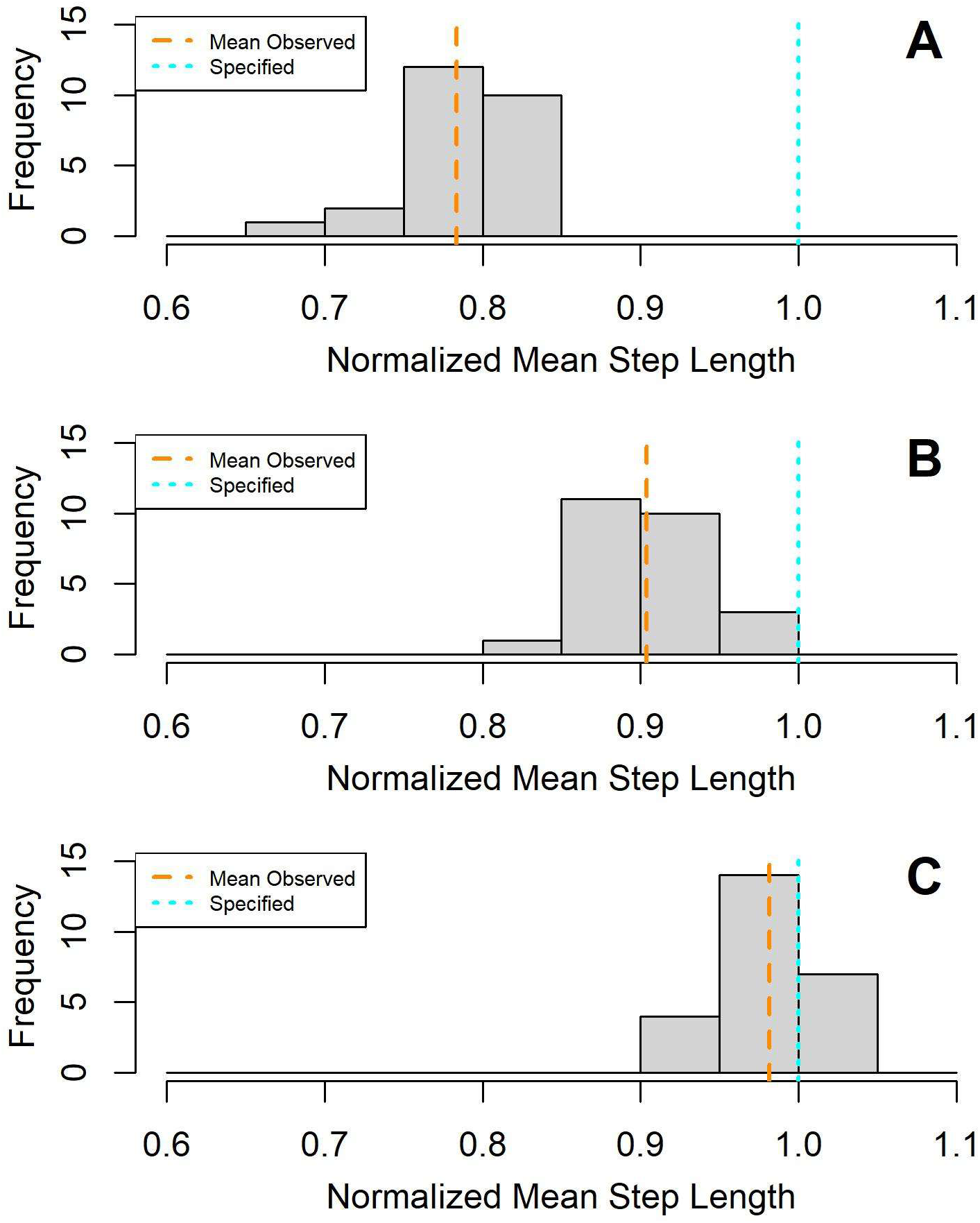
Normalized Mean Step Length by Smoothing Level, (A) W = 1, (B) W = 2, (C) W = 4

Researchers must estimate an animal’s mean step length in order to sample CSs to be paired with each OS. For instance, if researchers assume that step lengths follow an exponential distribution (as we have), the mean step length characterizes this distribution. In practice, though, researchers do not know the true mean step length of an animal. Instead, they may estimate the mean step from the animal’s OSs. Similarly, we fit an exponential distribution to the OSs and use this distribution to sample CSs. Because of the phenomenon of shorter steps described in the preceding paragraph, we typically obtain a distribution of step lengths with a mean step length that was less than what we had specified. As a result, our sampled CSs have mean step lengths that matched the OSs rather than what we had specified.

Unlike step lengths, relative turning angles do not appear to be affected by this phenomenon. Fig. S-16 shows the histogram across all 175 tracks. The mean of the mean observed relative turning angles (shown by the orange dashed line) differs from the teal dashed line by less than 0.002 radians.

**Fig. S-16.**
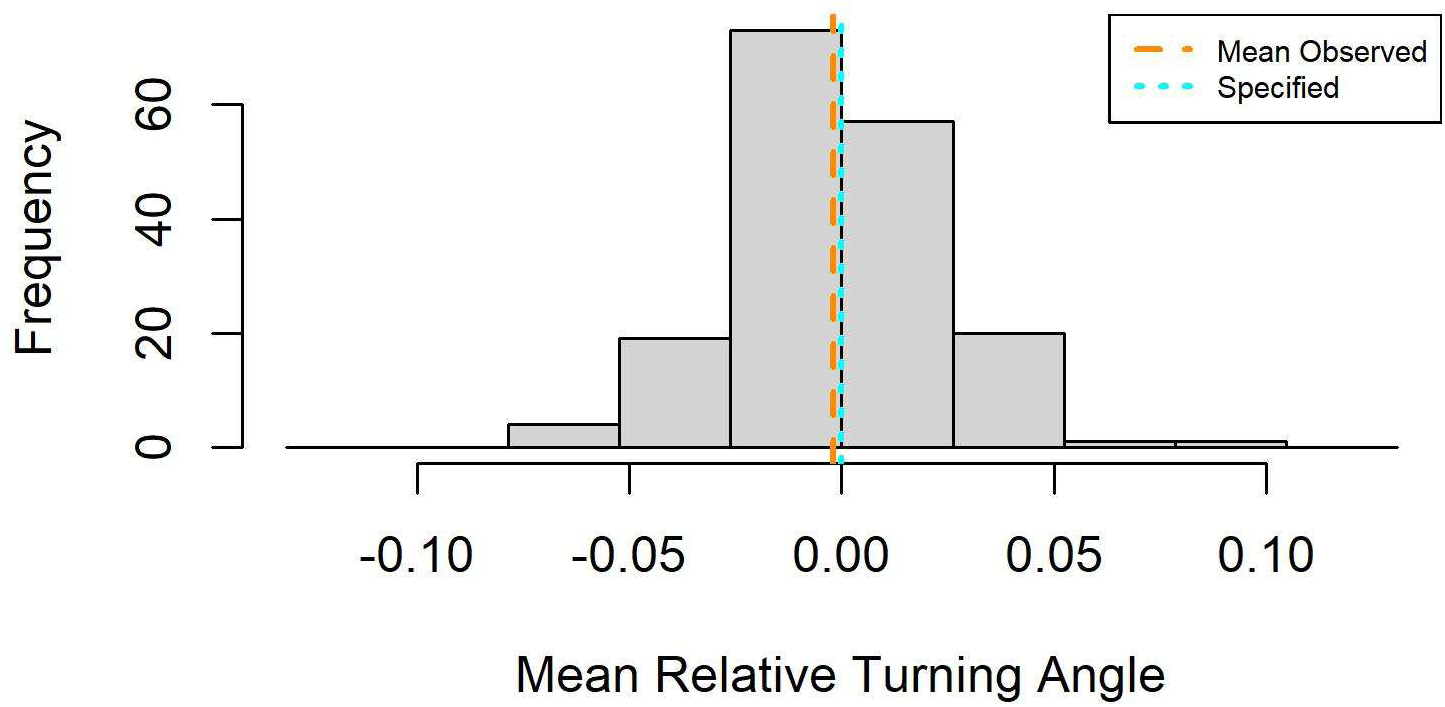
Mean Relative Turning Angles by Track

### Appendix F. Additional Results

Fig. S-17 and Fig. S-18 show additional results for tracks from the base class (with specified mean step length *μ_L_* = 4, smoothing window size *W* = 2, and track length *N* = 1000). Each plot corresponds to a given track and shows how the errors in estimated effects decrease as the number of CSs per OS increases. There are twelve plots in each of these two figures. In combination with Fig. 4 from the main paper, results from all 25 tracks in the base class are shown.

**Fig. S-17.**
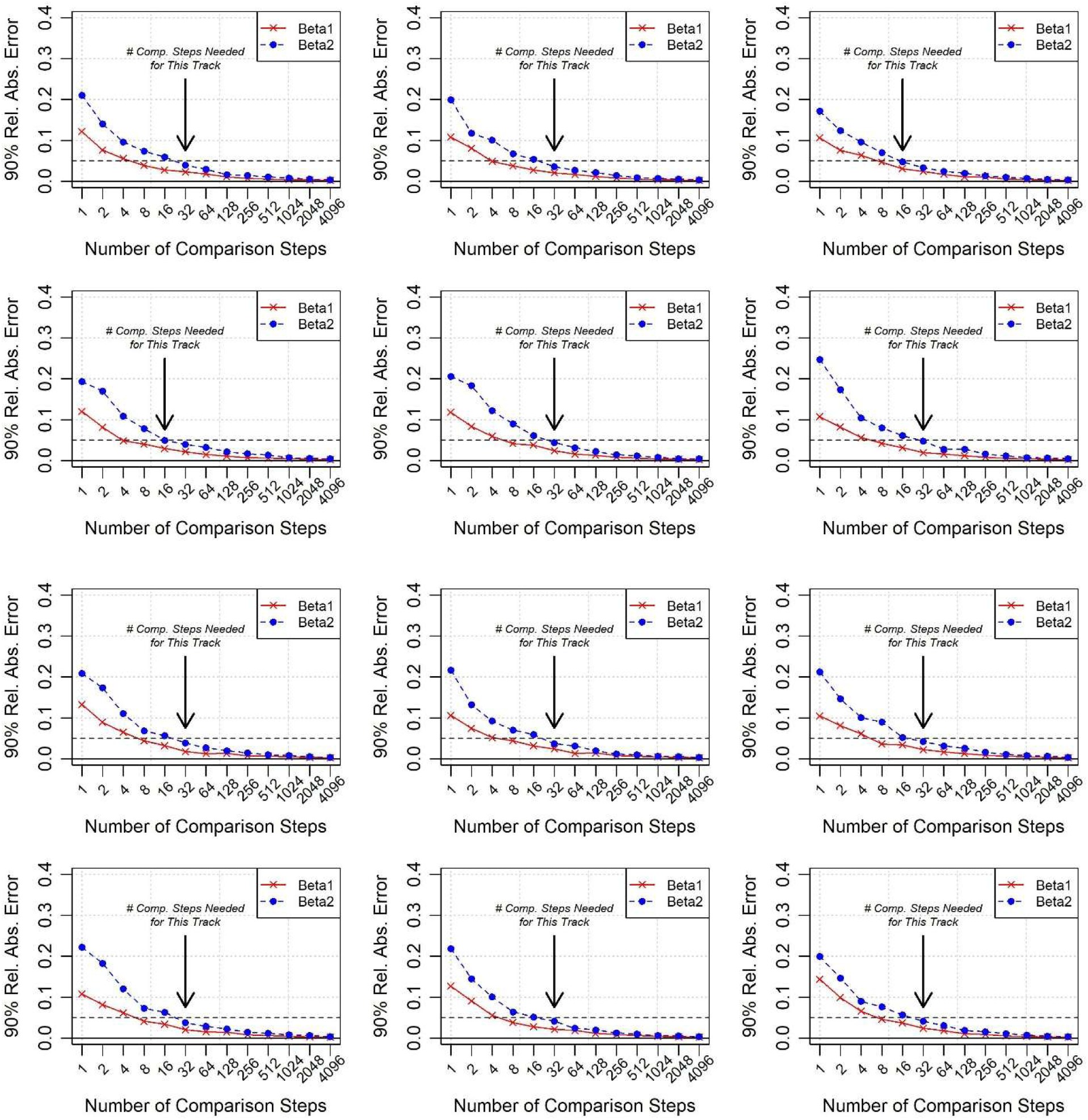
Number of Comparison Steps Needed for Individual Tracks in Base Class (Part 1)

**Fig. S-18.**
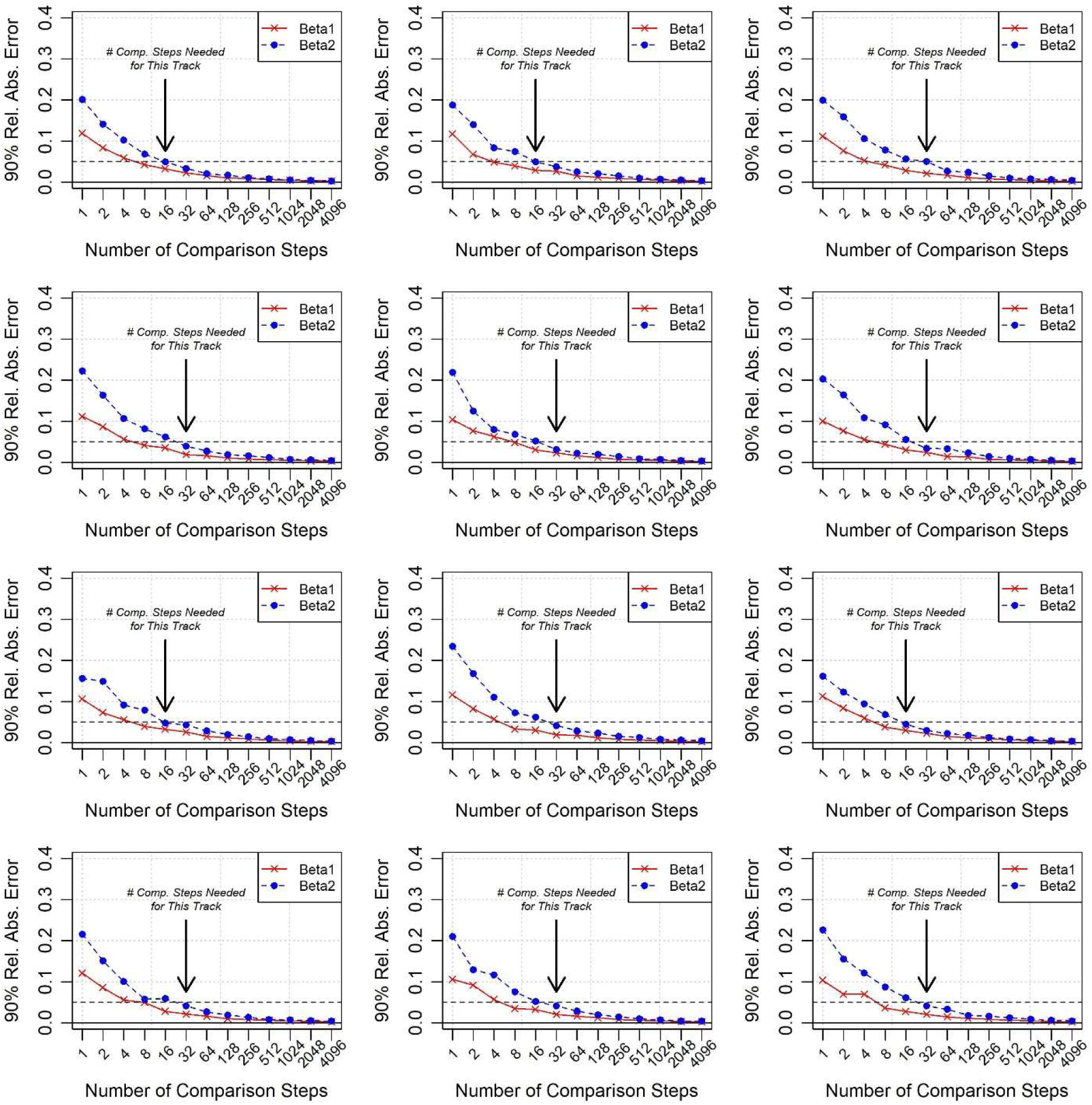
Number of Comparison Steps Needed for Individual Tracks in Base Class (Part 2)

The following figures compare the number of CSs needed for tracks with different characteristics. For each track, we used the criteria described in the main paper to determine the number of CSs needed. Fig. S-19 shows how the number of CSs needed changes with different amounts of landscape smoothing, while Fig. S-20 shows changes related to track length. Finally, Fig. S-21 shows changes in CSs needed as mean step length is varied. As discussed in the main paper, additional CSs are needed with smoother landscapes, shorter tracks, and shorter steps.

**Fig. S-19.**
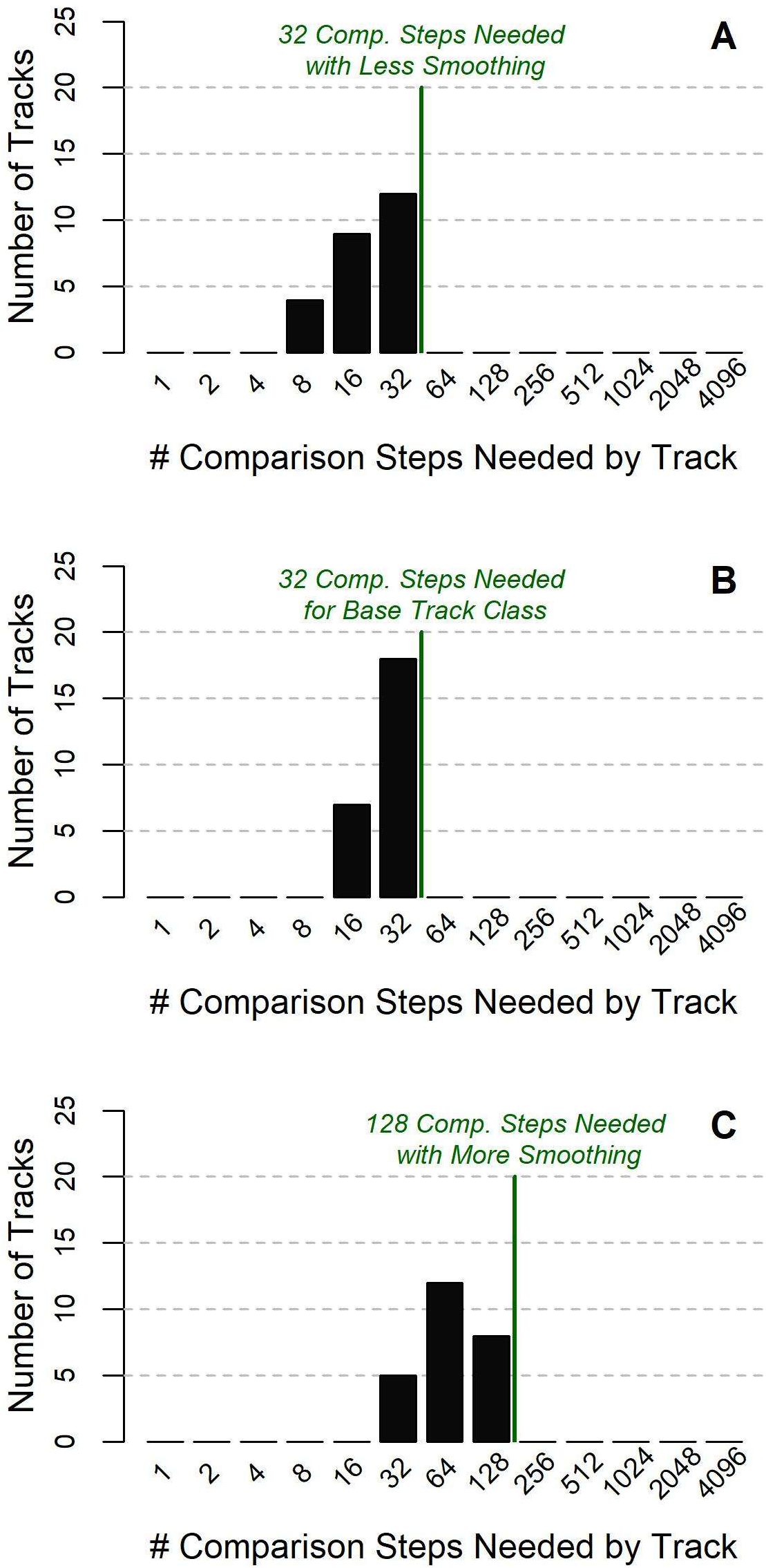
Distribution of Comparison Steps Needed for Tracks with (A) Less Smoothing, (B) Moderate Smoothing, (C) More Smoothing

**Fig. S-20.**
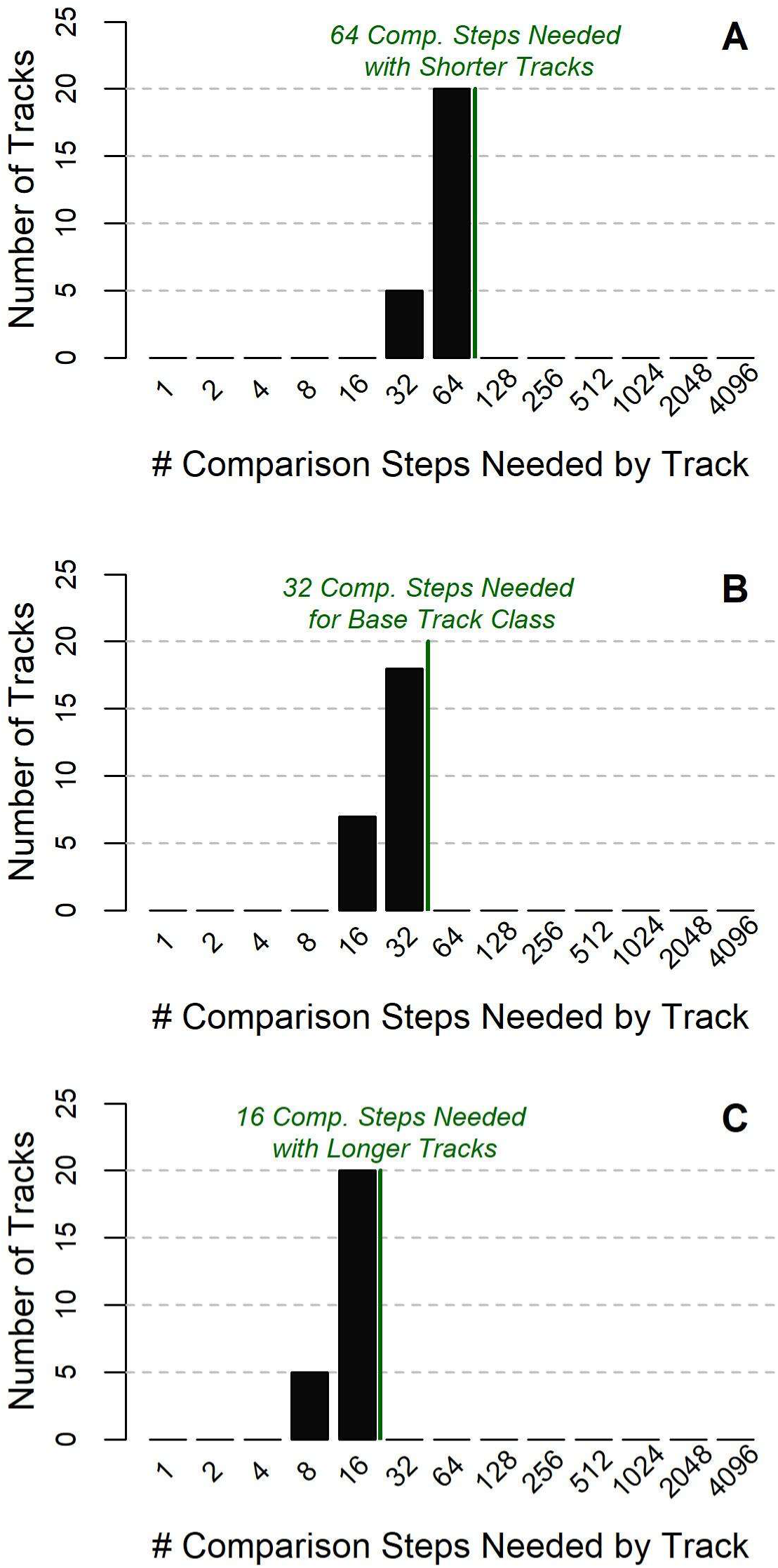
Distribution of Comparison Steps Needed for Tracks with (A) Shorter Tracks, (B) Moderate Track Lengths, (C) Longer Tracks

**Fig. S-21.**
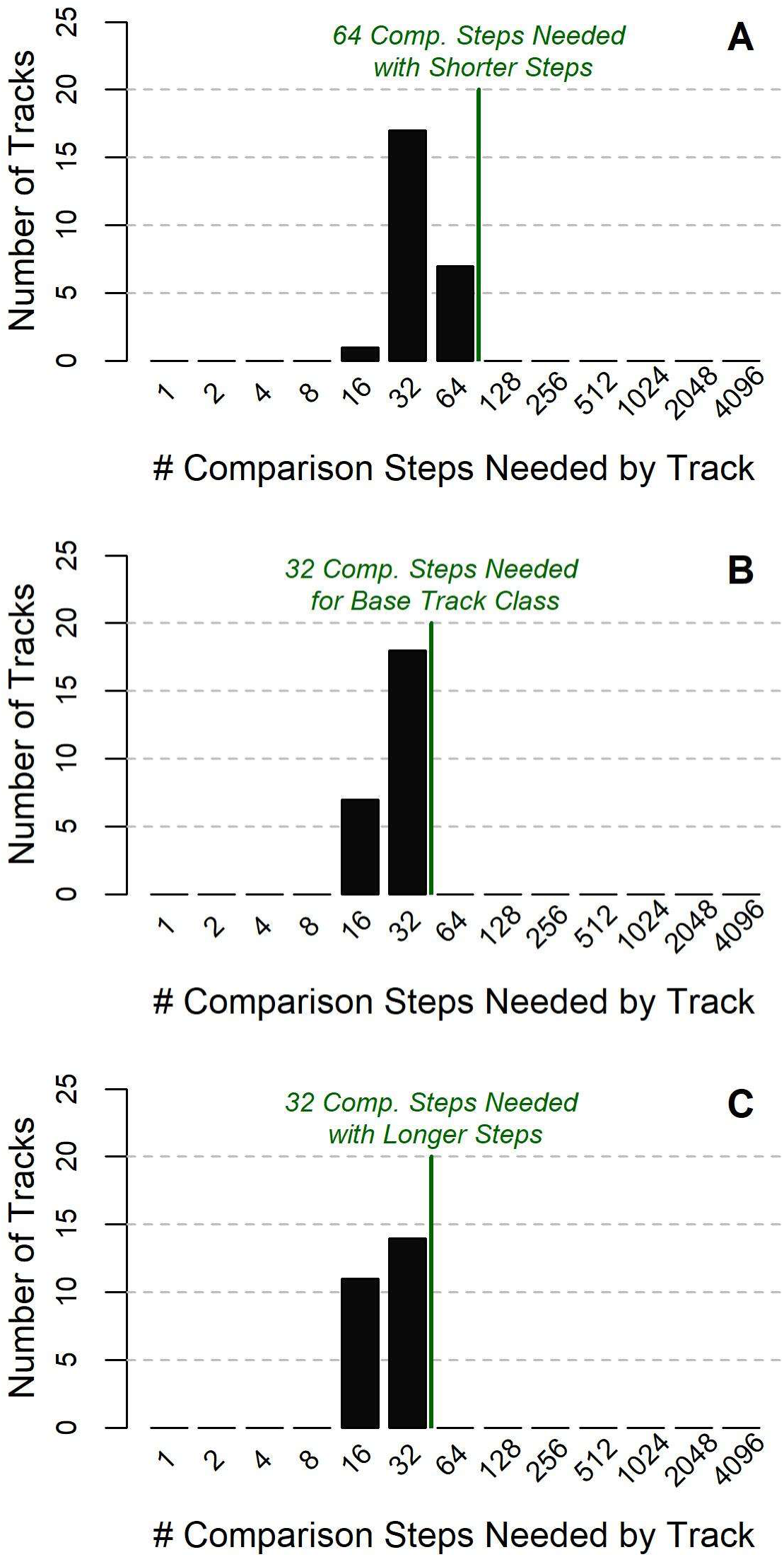
Distribution of Comparison Steps Needed for Tracks with (A) Shorter Steps, (B) Moderate Step Lengths, (C) Longer Steps

We primarily used 5% (*ε_1_* = 0.05) as the RAE threshold for determining when the estimates for a given parameter had “converged” (see Section 3 in the main paper). In particular, we required that the RAEs for 90% of all replicates with a given number of CSs were below 5%. Aside from its acceptance as a popular convention, the selection of 5% for this threshold is arbitrary. To assess the sensitivity of our conclusions to this selection, we repeated our analysis with *ε_1_* = 0.01. Fig. S-22 shows a distribution of the number of CSs needed for tracks in the base class using this lower threshold value. As expected, requiring a lower threshold (which is more demanding) results in larger numbers of CSs needed.

**Fig. S-22.**
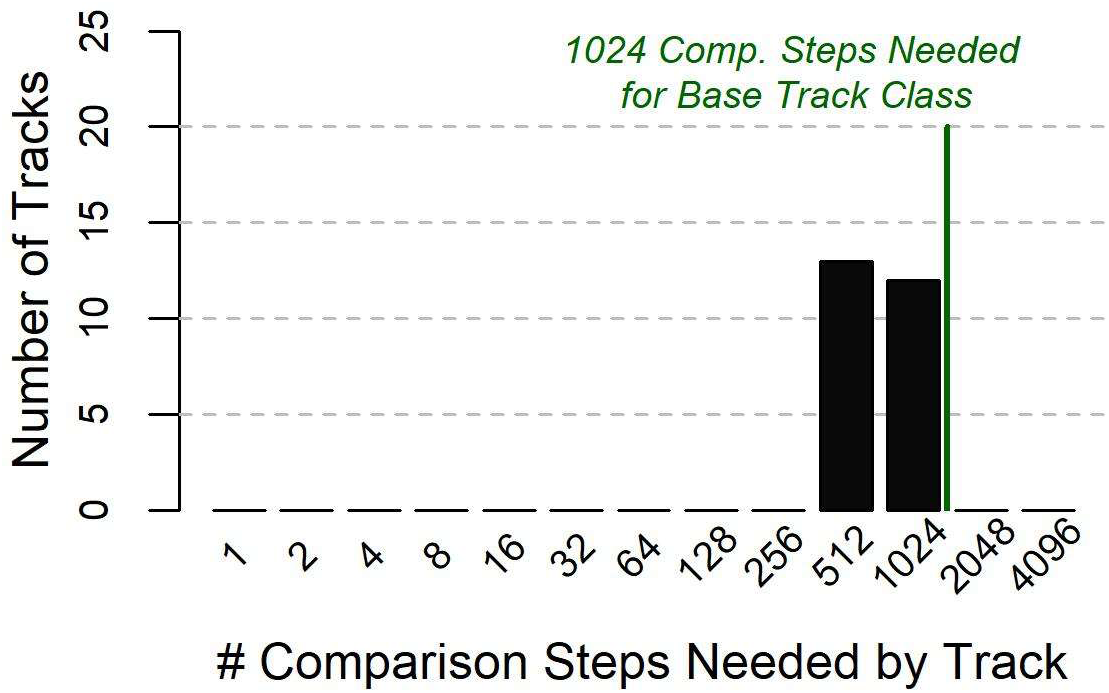
Distribution of Comparison Steps Needed by Track with Lower Relative Absolute Error Threshold

Within our analysis, we also primarily used RAE as the metric by which convergence was determined. This metric directly compares estimates using fewer CSs with those estimated with many CSs (“true simulated effect” β^*^_*i*_). We might instead track the variation of the estimated effects while the number of CSs is held constant, e.g., with a relative standard deviation (RSD). This variation is due to the random sampling of the CSs. For a given covariate *z*_i_ with the standard deviation of its estimates as *σ*_i_, we compute *RSD_i_* = σ_*i*_/β^*^_*i*_. Fig. S-23 shows how RSD values changes as the number of CSs increase for an example track.

**Fig. S-23.**
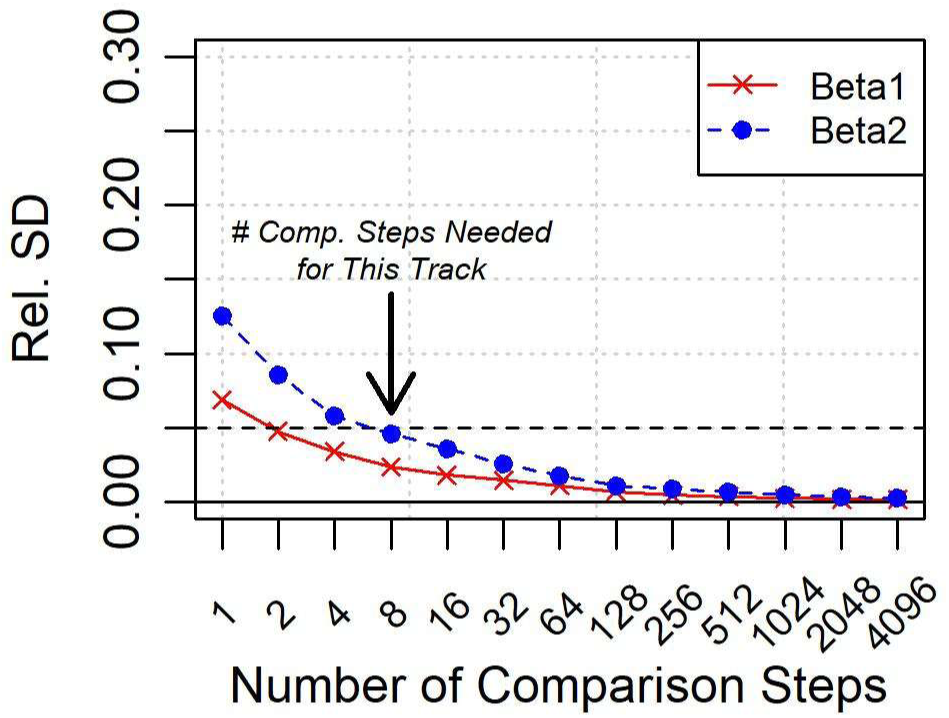
Number of Comparison Steps Needed for An Example Track, According to Relative Standard Deviation

As with RAE, we can specify the threshold *ε_1_* for determining convergence when assessing RSD for each number of CSs. Fig. S-24 shows the number of CSs needed for each track in the base class when *ε_1_* = 0.05. Likewise, Fig. S-25 shows this distribution when *ε_1_* = 0.01.

**Fig. S-24.**
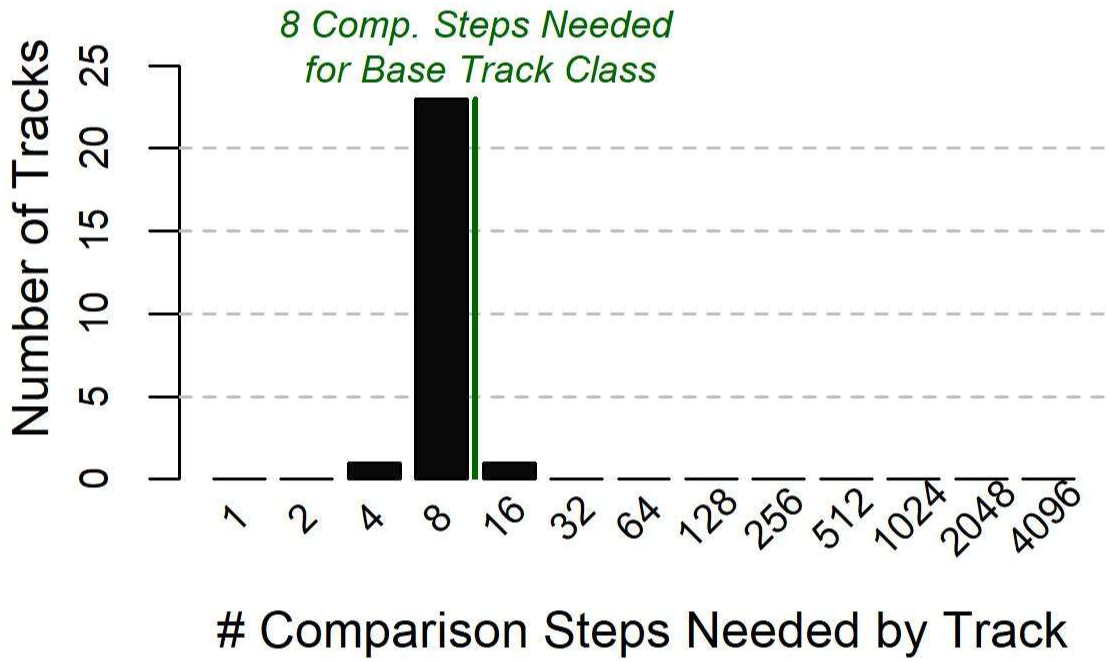
Distribution of Comparison Steps Needed, According to Relative Standard Deviation

**Fig. S-25.**
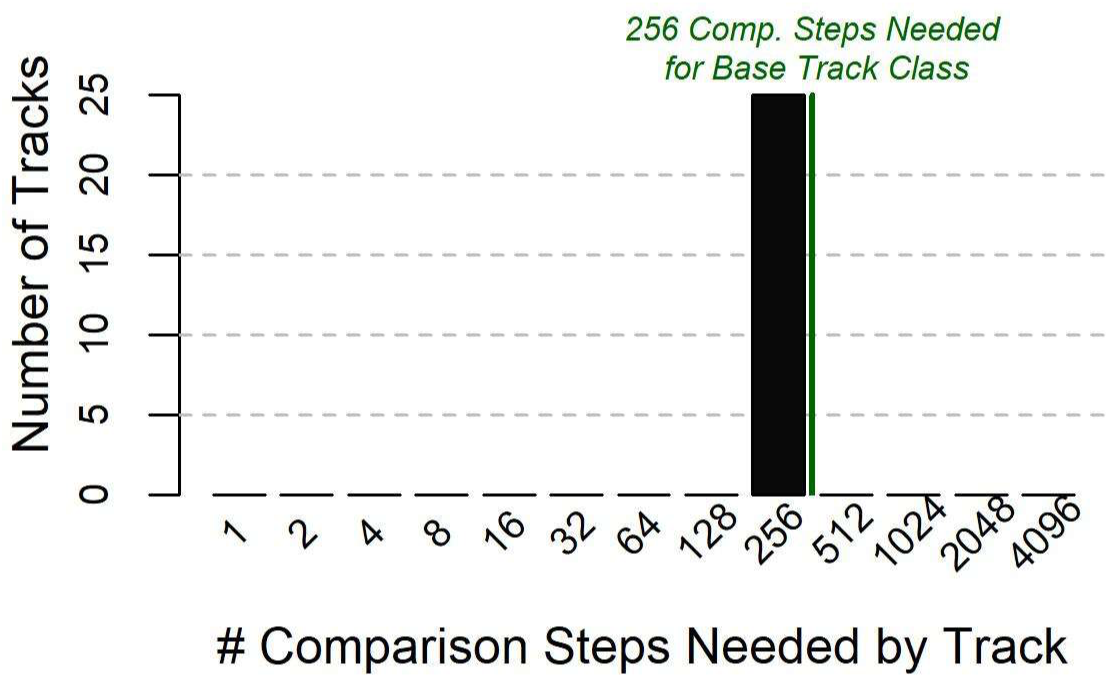
Distribution of Comparison Steps Needed, According to Relative Standard Deviation and with Lower Convergence Threshold

To examine these trends with greater resolution, we also computed the mean 90^th^ percentile RAE across all numbers of CSs and all tracks within a given class of tracks (Table S-5). These values do not directly assess how many CSs are needed for individual tracks. However, they enable even comparison of the influence of landscape and track characteristics on effect estimation accuracy. Moreover, since errors in estimated effects guide how many CSs are needed for a given track, this metric assesses the interaction between the number of CSs needed and landscape or track characteristics. These values confirm the trends observed in Table 2. Greater effect estimation errors arise for tracks with relatively greater landscape smoothing, shorter mean step lengths, or shorter track lengths.

**Table S-5:**
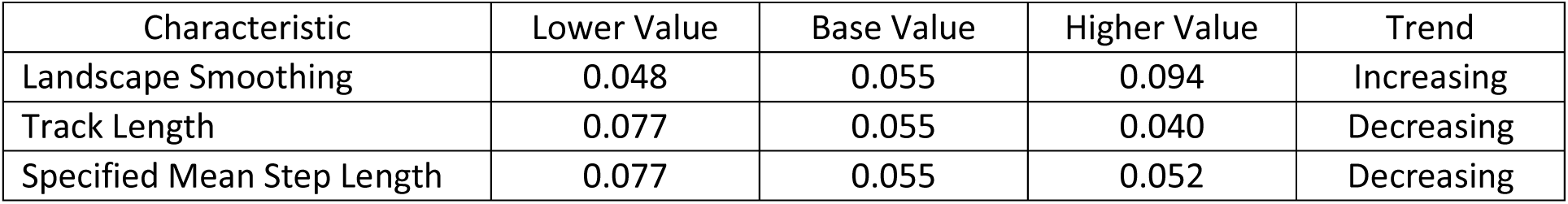
Mean Relative Absolute Error for Each Class of Tracks.

| Characteristic | Lower Value | Base Value | Higher Value | Trend |
| --- | --- | --- | --- | --- |
| Landscape Smoothing | 0.048 | 0.055 | 0.094 | Increasing |
| Track Length | 0.077 | 0.055 | 0.040 | Decreasing |
| Specified Mean Step Length | 0.077 | 0.055 | 0.052 | Decreasing |

### Appendix G. Description of Available Data and Code

The repository posted at https://github.com/StatsForAnimalMovement/InsightsForSSFs/contains 175 R workspace files (*.RData*) – one for each of the 25 tracks within each of the 7 track classes discussed in this paper. Each file contains several data objects:

- *Track* (data frame): The simulated track is expressed in terms of x- and y-coordinates. For subsequent use as inputs in step selection functions, step length, absolute angle (computed relative to movement in the positive direction along the x-axis), and relative angle (computed relative to the previous step) are also included.
- *ParamList* (data frame): The parameters used to simulate the track have been recorded in this data frame.

`o` *LandscapeNum* = index of the landscape on which the track has been simulated (varies from 1 to 5)
`o` *TrackNum* = index of tracks within each class and on each landscape (varies from 1 to 5)
`o` *Beta1* = the specified effect for the first covariate (set to 4 for all tracks in this study)
`o` *Beta2* = the specified effect for the second covariate (set to 2 for all tracks in this study)
`o` *TrackLength* = number of steps in the track (set to 500, 1000, or 2000 in this study)
`o` *CorWindow* = smoothing window size that specifies the width of the window used to smooth each grid within the covariate landscape (set to 1, 2, or 4 in this study); the width of the window is equal to 2\**CorWindow* + 1
`o` *StepLengthParam* = reciprocal of the mean step length for the track (set to 1/8, 1/4, or 1/2 in this study)
`o` *TurningAngleKappa* = parameter that determines the width of the peak of the von Mises distribution from which the relative turning angles of candidate steps are drawn (set to 3 for all tracks in this study); higher values yield tighter peaks
- *ResultsByModel* (data frame): The fitted effects for each model from each replicate with different numbers of comparison steps have been recorded in this data frame. There is one entry for each model.

`o` *LandscapeNum* = index of the landscape on which the track has been simulated (varies from 1 to 5)
`o` *TrackNum* = index of tracks within each class and on each landscape (varies from 1 to 5)
`o` *NumCompSteps* = number of comparison steps paired with each observed step
`o` *Round* = replicate number for a given track and a given number of comparison steps
`o` *PointEstimate1* = point estimate for the first covariate’s effect (from a model fitted without the noise covariate)
`o` *StandardError1* = standard error for the first covariate’s effect (from a model fitted without the noise covariate)
`o` *PointEstimate2* = point estimate for the first covariate’s effect (from a model fitted without the noise covariate)
`o` *StandardError2* = standard error for the first covariate’s effect (from a model fitted without the noise covariate)
`o` *AIC2* = AIC of a model fitted with only the two true covariates, i.e., without the noise covariate
`o` *AIC3* = AIC of a model fitted with the two true covariates that influenced the track simulation, as well as the noise covariate

Using the *ResultsByModel* data frame, the enclosed code (*findNumCompStepsNeeded.R*) allows the number of CSs needed for each track to be determined.

## List of Abbreviations

CS: Comparison step
OS: Observed step
RAE: Relative absolute error
RSD: Relative standard deviation
SSF: Step selection function

